# The Paradox of the Plankton: Coexistence of Structured Microbial Communities

**DOI:** 10.1101/2021.09.13.460068

**Authors:** Alberto Scarampi

## Abstract

In the framework of resource-competition models, it has been argued that the number of species stably coexisting in an ecosystem cannot exceed the number of shared resources. However, plankton seems to be an exception of this so-called “competitive-exclusion principle”. In planktic ecosystems, a large number of different species stably coexist in an environment with limited resources. This contradiction between theoretical expectations and empirical observations is often referred to as “The Paradox of the Plankton”. This project aims to investigate biophysical models that can account for the large biodiversity observed in real ecosystems in order to resolve this paradox. A model is proposed that combines classical resource competition models, metabolic trade-offs and stochastic ecosystem assembly. Simulations of the model match empirical observations, while relaxing some unrealistic assumptions from previous models.

**Paradox:** from Greek *para*: “distinct from”, and *doxa*: opinion. Sainsbury (1995) defines a paradox as “an apparently unacceptable conclusion derived by apparently acceptable reasoning from apparently acceptable premises”. Paradoxes are useful research tools as they suggest logical inconsistencies. In order to spot the flaw, the validity of all the premises has to be carefully assessed.

**Plankton:** refers to the collection of organisms that spend part or all of their lives in suspension in water (Reynolds 2006). Plankton, or *plankters*, are “organisms that have velocities significantly smaller than oceanic currents and thus are considered to travel with the water parcel they occupy” (Lombard et al. 2019). *Phytoplankters* refer to the members of the plankton that perform photosynthesis.

## 1 Introduction

Our planet is home to an unthinkable amount of life: up to a trillion (10^12^) species (Mora et al. 2011; Lombard et al. 2019) thrive on Earth, a number which approximates to the estimated number of galaxies in the universe (Conselice et al. 2016). This leads to the question: how can there be so much biodiversity on Earth? Ecologists often explain this in terms of ecosystems. In nature, different species assemble into stable communities, coexisting in the same environment. These “societies of species” emerge from the interactions between individual organisms that are competing for the same finite amount of space and resources. How such coexistence emerges from collectives of competing individuals is however still not clear. These ecosystems are not only able to robustly adapt to a wide range of environments, ranging from the ocean to the human gut, but also actively modify their surroundings, influencing large-scale planetary cycles (Falkowski et al. 1998), and are of great economical interest for human societies (Brenner et al. 2008; Turnbaugh et al. 2006). Given the the future regimes of unpredictable climate perturbations and the aims of a more circular, sustainable bioeconomy, it is a vital challenge to understand the forces that drive ecosystems and predict their biodiversity.

### 1.1 The Paradox

An interesting case to investigate the forces that drive ecosystems is offered by plankton (see page 3 for definition). Hutchinson (1961) published a paper entitled “The Paradox of the Plankton”. He pointed out that “the problem that is presented by the plankton is essentially how it is possible for a number of species to coexist in a relatively unstructured environment all competing for the same sorts of materials”. Plankton seems to disobey the principle of competitive-exclusion, the classical mathematical framework commonly used to interpret the dynamics of competitive ecosystems. Although there is nothing absurd about plankton, its case follows the definition of what is generally known as a paradox: “an apparently unacceptable conclusion derived by apparently acceptable reasoning from apparently acceptable premises” (Sainsbury 1995). In the paradox of the plankton, the unacceptable conclusion derives from the principle of competitive-exclusion, the reasoning is the mathematical framework of competitive population dynamics, and the premises are the mathematical assumptions used to derive the model. As paradoxes might arise from false statements, it is useful to explain the principle that plankton seems to contradict. The principle of competitive exclusion postulates that the number of species (*m*) coexisting in a “well-mixed” ecosystem, at steady state, cannot exceed the number of shared limiting resources (*p*) (Hardin 1960). The mathematical premises of the principle are attributed to Volterra (1928), who realised that coexistence of two species (*σ*_1_, *σ*_2_) limited by the same shared resource, *c*_*i*_ is impossible (Armstrong & McGehee 1980). Assuming that organisms increase or decrease in a continuous way, Volterra (1928) describes the dynamics of *σ*_1_ and *σ*_2_, both consuming *c*_*i*_, using the ordinary differential equations (ODEs) below:

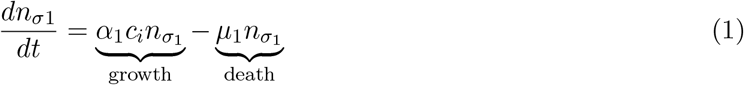

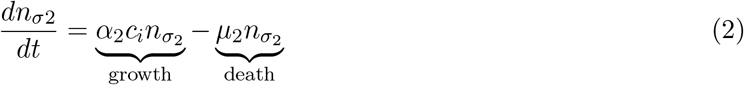

The change in population size of the two species 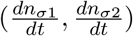 is determined by the difference between the growth rates and death rates of each species. Every species grows proportionally to its rate of nutrient uptake (*α*_*i*_) and dies linearly according to the death rate *δ*. Both species compete for the same resource in a “well-mixed” environment. Therefore, *c*_*i*_ can be written as the difference between its maximum nutrient concentration (in the absence of any species) and the amount consumed by the two species, according to Equation 3.

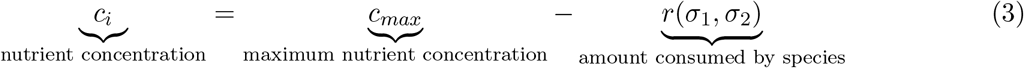

Where *r*(*σ*_1_, *σ*_2_) is a function describing the uptake of resources by the two species. After substituting Equation 3 into Equation 1 and 2, the latter two can be rewritten as below:

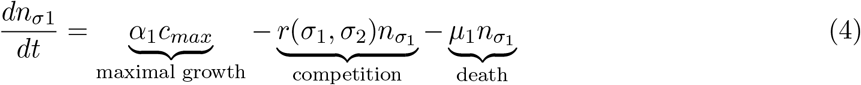

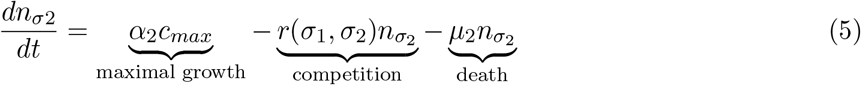

Setting *α*_1_*c*_*max*_ − *μ*_1_ = *ϵ*_1_ and *α*_2_*c*_*max*_ − *μ*_2_ = *ϵ*_2_ results in

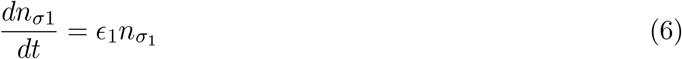

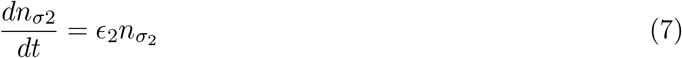

Volterra (1928) showed that as time approaches infinity, the species with the largest rate of increase (*ϵ*_*i*_), will approach a finite nonzero density, whereas the second one will inevitably face extinction. The principle has been later generalised to the case of *m* < *p* species (Armstrong & McGehee 1980; Rescigno & Richardson 1965). Inevitably, the species with the largest *ϵ*, will drive the latter to extinction (Hardin 1960). However, as illustrated in Figure 1, planktic ecosystems appear to disobey the postulates of competitive exclusion. While limited by mainly 3 resources (Hutchinson 1961), more than 100 different phytoplankters can coexist in a litre of seawater (Davies et al. 2016).

**Figure 1:**
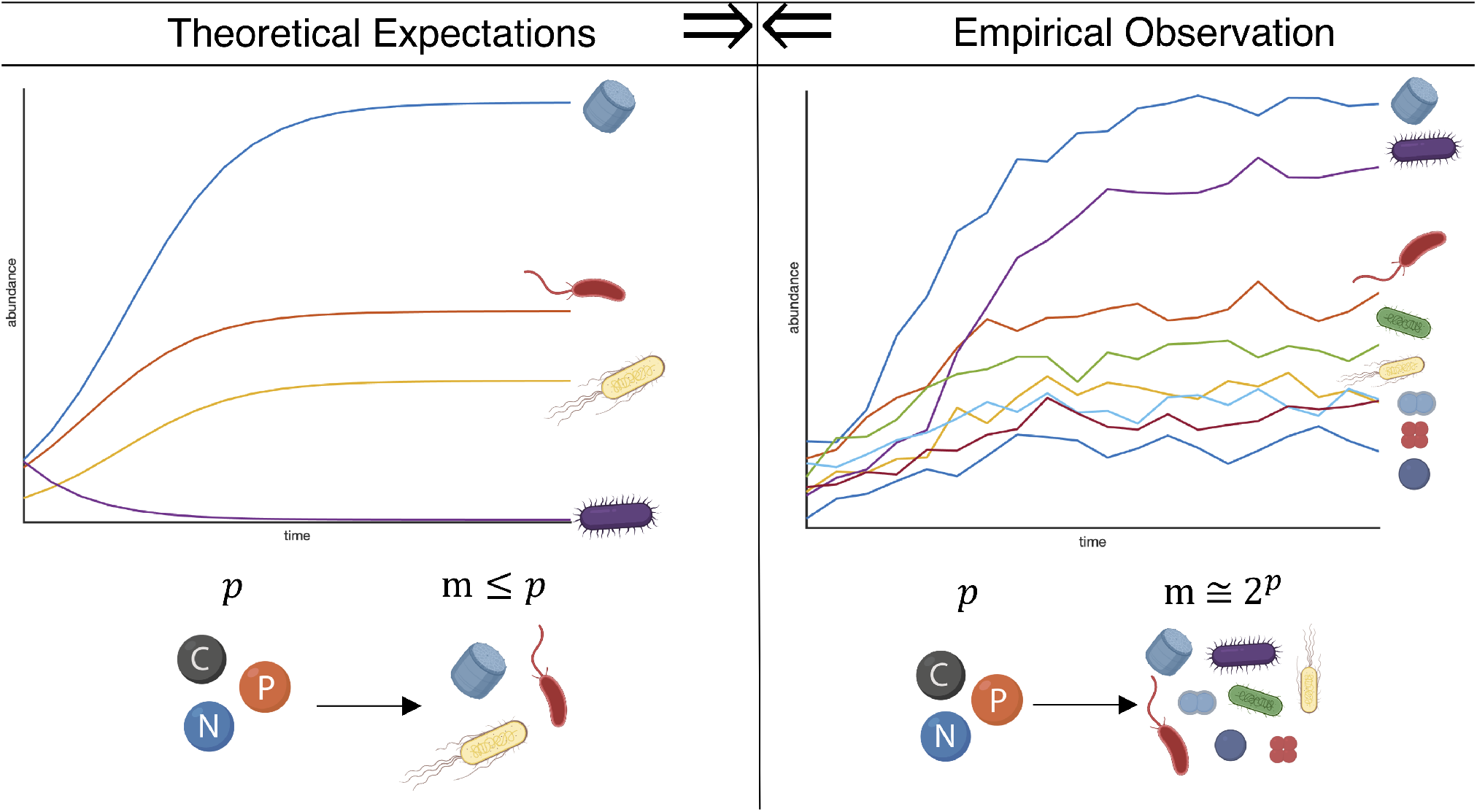
Phytoplankters (illustrated as the colourful cells) are generally assumed to be limited by *p* = 3 resources: carbon (the black sphere) nitrogen (blue) and phosphorus (orange). Resource competition models (on the left) predict that the number of coexisting species (*m*) cannot exceed the number of resources (*p*), hence *m* ≤ *p*. On the other hand, observations of real ecosystems (right hand side) reveal exponential relationship between the number of resources and the the number of coexistence species, *m* ≈ 2^*p*^. This contradiction (⇒⇐) between theoretical models for ecosystem growth and experimental observation is known as “The Paradox of the Plankton”.

### 1.2 The Assumptions

Scheffer et al. (2003) offers an overview of the main proposed resolutions to the paradox: nonequilibrium dynamics, non-linear dynamics and spatial heterogeneity. These all refute some of the assumptions made by Volterra (1928). This thesis aims to identify flaws in the premises leading to the paradox. Therefore it is useful to clearly state these suppositions.

I. **Steady State Assumption:** The principle of competitive exclusion derives from assuming that species reach a steady-state. However, it is hard to find scientific evidence of such equilibrium. In fact, plankton growth has been shown to be chaotic (Benincá et al. 2008).
II. **Limiting Resources Assumption:** Volterra (1928) assumes that species growth only depends on shared limiting resources. In reality, ecosystems are limited by factors other than resources, such as predation Laan & de Polavieja (2018), self-limiting toxin production (Geyrhofer & Brenner 2018) and trade-offs between different metabolic abilities (Posfai et al. 2017).
III. **“Well-Mixed” Assumption:** The principle of competitive-exclusion derives from the assumption of spatial homogeneity in the ecosystem.

Whereas models that relax the first two assumptions are ubiquitously found in the literature (Bonachela et al. 2011; Huisman et al. 2001; Posfai et al. 2017), spatial heterogeneity is an often talked about but rarely modelled resolution to the paradox. Plankton has historically been assumed homogeneous as it is composed of microorganism floating on turbulent water. Indeed, Hutchinson (1961) himself found it “hard to believe that in turbulent open water many physical opportunities for niche diversification exist”. However, recent scientific evidence demonstrates that non “well-mixed” distributions of species and resources across different spatial scales is a common scenario in natural ecosystems (Malviya et al. 2016). Indeed, biophysical processes affecting plankton ecology span a range of scales (Prairie et al. 2012), the larger the scale the more significant the heterogeneities. At the micro-scale, the scale of micrometers and less, dissolved organic matter secreted by microorganisms is spatially partitioned by molecular size and attracts bacteria in discrete patches (Smriga et al. 2016). At the meso-scale, the scale in between micrometers and meters, phytoplankton cells can be spatial separated in turbulent vortex structures (Bracco et al. 2000, Durham et al. 2013). Using diffusion-reaction equations, Lindemann et al. (2017) showed that turbulence produced by eddies can separate cells by their sizes, which determine their sinking rates. At the macro-scale, the scale of meters and more, an entangled network of environmental forces (e.g. winds, oceanic currents), contribute to create heterogeneous distributions of phytoplankton communities, with sizes and structures varying across different oceanic regions (Acevedo-Trejos et al. 2015). Furthermore, Guirey et al. (2010) analytically demonstrated that large-scale diffusion, far from disrupting plankton patchiness, actually results in the emergence of spatial structures.

### 1.3 Aims

This thesis aims to clarify the paradoxical situation of the plankton, by investigating a biophysical model that combines generalised Lotka-Volterra equations with trade-off consumer-resource dynamics. By systematically evaluating the resulting dynamics of the proposed model with a simple stochastic computer simulation, the aim is to answer the following questions:

- How does coexistence emerge from collectives of competing individuals?
- Can spatial heterogeneity account for the high biodiversity observed in real ecosystems ?
- What factors/assumptions lead to the “Paradox of the Plankton”?

## 2 Methods - Modelling Framework

To address the questions above, a simple biophysical model will be investigated and simulated *in silico*. The model combines trait-based modelling (Litchman & Klausmeier 2008), generalised Lotka-Volterra growth equations (Gonze et al. 2018) and stochastic ecosystem assembly (Geyrhofer & Brenner 2018). Since biophysical processes affecting plankton ecology span a range of scales (Prairie et al. 2012), a multi-scale framework based around the concept of metacommunity is employed (Leibold et al. 2004). The ecosystem here is thus modelled as a discrete set of sub-ecosystems connected by spatial flows of energy, materials and organisms across ecosystem boundaries (Loreau et al. 2003). Different scales of ecosystems are simulated hierarchically: micro-scale interactions between species and nutrients, mesoscale homogeneous communities (hereafter referred to as demes) and macro-scale metacommunities (communities of communities). The different scales of the ecosystem are modelled as classes using object oriented-programming in MATLAB. Classes are described by their properties and functions that can be applied to them. This approach makes the simulator modular and easily extendable for further investigations.

### 2.1 Modelling Cells: Species

At the micro-scale level, the ecosystem is populated by cells interacting with the surrounding environment and nutrients.

#### Definition

A species *σ* is defined as an instance of the `Species` class, and is constructed by inputting a name (*σ*_*i*_), an array of traits 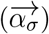 and the size of its population (*n*_*σ*_).

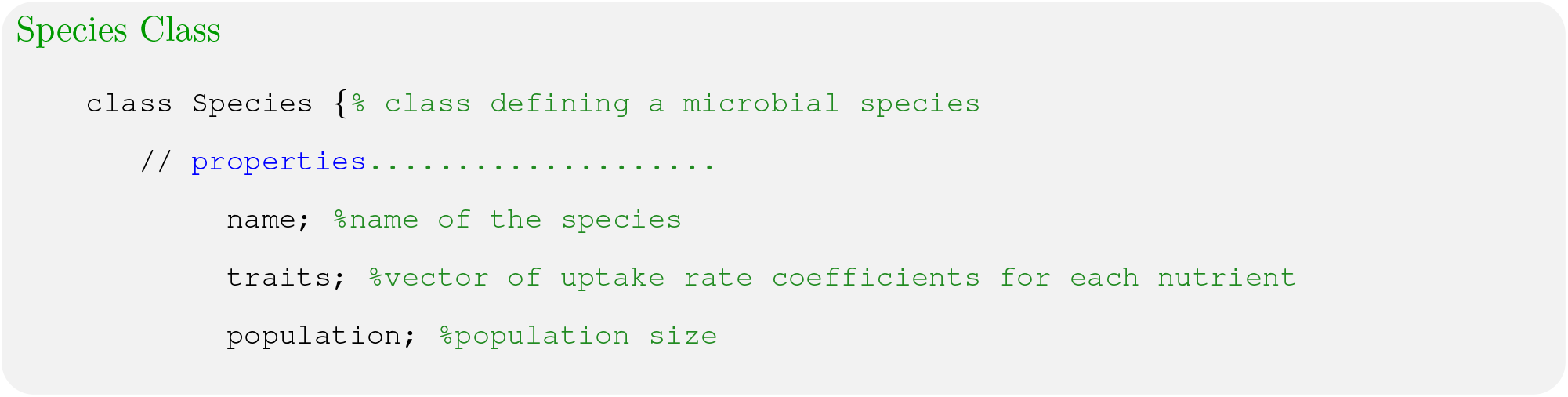

#### Assumptions

I. **Trait-Based Modelling:** Every species is defined by a vector of traits 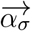. Traits are the quantifiable characteristics that distinguish each species from another. Modelled traits often include cell size, nutrient uptake rates and ecological roles (Litchman & Klausmeier 2008). As this model investigates microbial coexistence within an ecosystem with spatial heterogeneity in the nutrient supply, nutrient uptake rates are the only traits taken into account. Every species is thus specified by its coefficients of rate of consumption for each nutrient 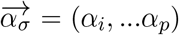. Biologically, *α_σi_* correlates to the amount of enzymes that each species invests to uptake/metabolise nutrient *i* (Posfai et al. 2017).
II. **Metabolic Trade-Offs:** Species are initialised with uniformly distributed random traits (*α*_*σi*_), to reflect different evolved nutrient consumption abilities. Different species can express different enzymes to process different nutrients. However, to account for trade-offs in resource utilisation, all organisms have the same fixed enzyme budget *E*:

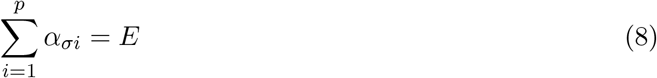 Because the sum of each species traits is a constant, the species can be visualised as points intersecting the plane *E* = *α*_1_ + *α*_2_ + *α*_3_ in the space of resource utilisation (Figure 3).
III. **Monod Uptake Rate:** The growth rate of each species depends on the availability of each nutrient. It is assumed that the internal (cytosolic) concentration is a function of the external nutrient concentration and proportional to the rate at which that nutrient is uptaken, *r_i_*(*c*_*i*_). The nutrient uptake function, *r*_*i*_, is described by Equation 9, which increases monotonically and saturates at high concentration of resource.

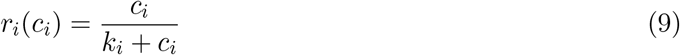 Where *k*_*i*_ is the half-saturation constant, similar to the Michaelis-Menten constant in enzyme kinetics (Johnson & Goody 2011).

**Figure 2:**
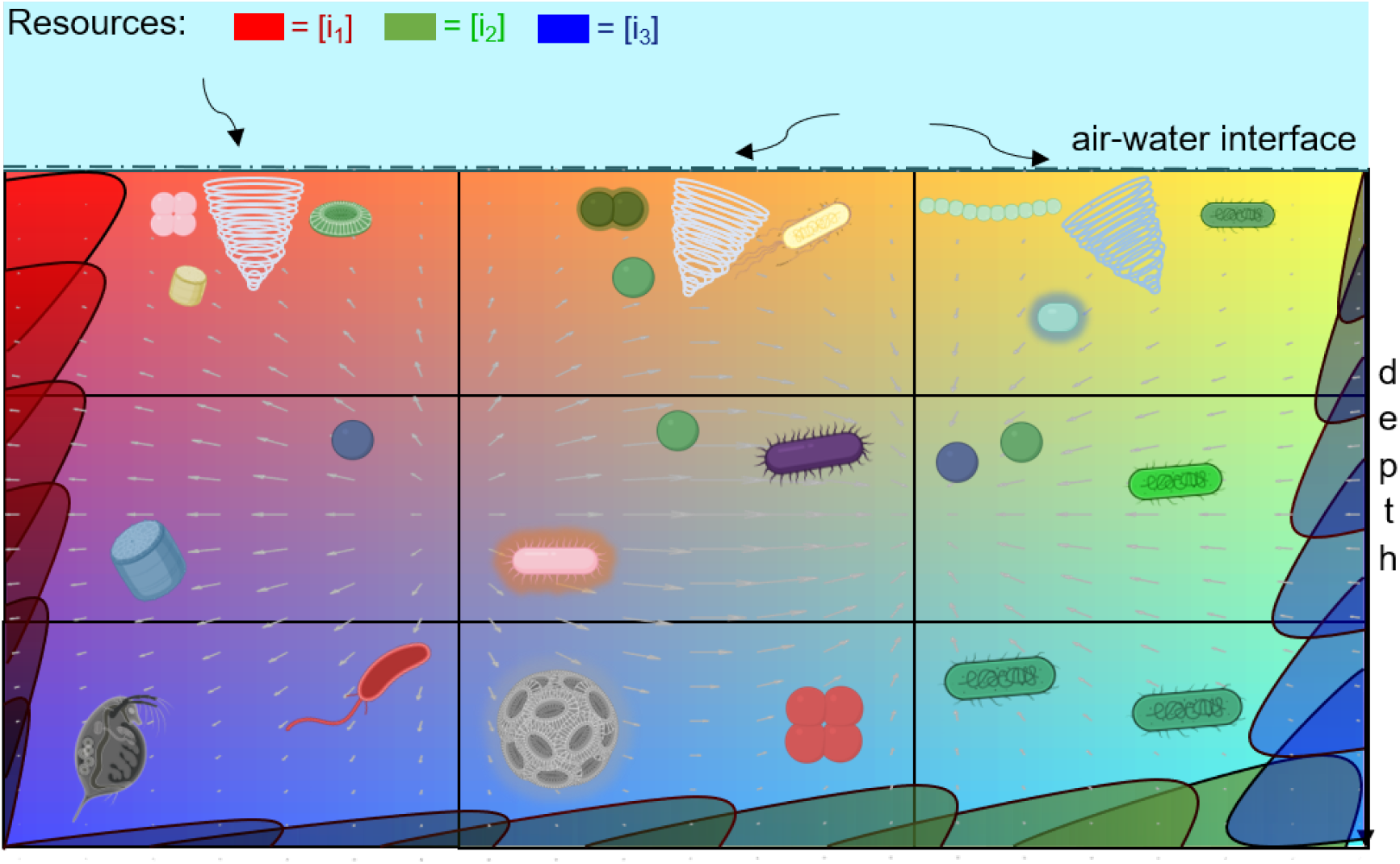
Evidence suggests that plankton is not a “well-mixed” ecosystem. Ocean mesoscale eddies (the white spirals) generated by air flows (curved black arrows) at the air-water interface can segregate different species in different compartments. Turbulence in water (the white vector field) can separate different species with different sinking rates at different depths in the water column. Resources is represented as a colour. Resource 1 = red, Resource 2 = green, Resource 3 = blue. The coloured gradients represent increasing concentrations. Resource 1 increases in concentration (increasing red) at decreasing depths. Thus it could represent inorganic carbon. Resource 2 increases in concentration from left to right. Thus it could represent phosphorus (if the left border is assumed to be the open ocean and the right border a hypothetical coast). Resource 3 increases at increasing depths. Thus it could represent nitrogen (which is fixed by diazotrophic cyanobacteria and sinks towards the bottom of the ocean when these are eaten by predators).

**Figure 3:**
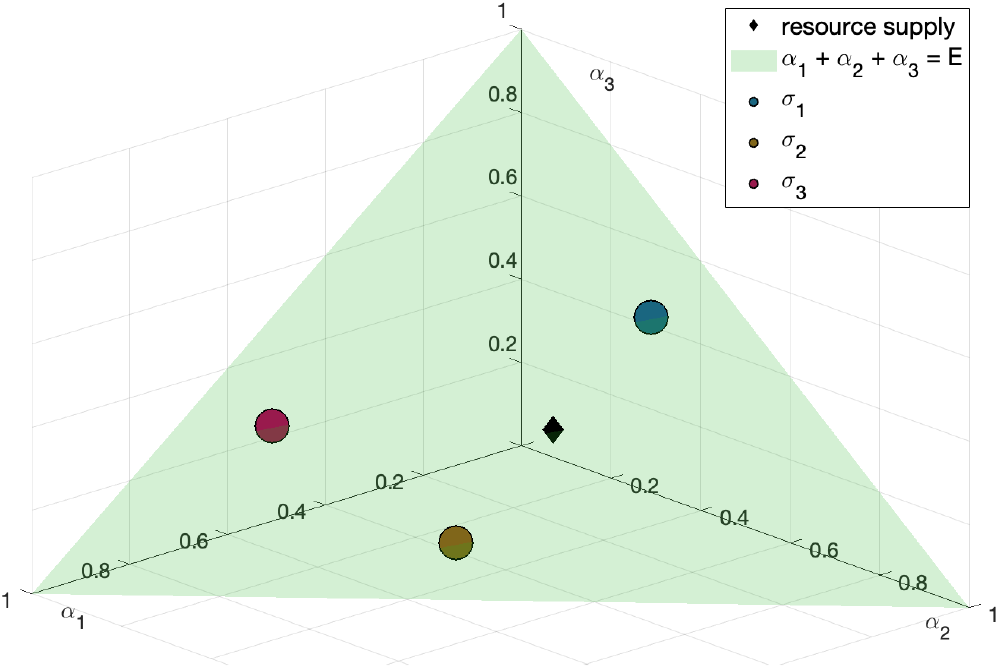
Every species occupies a niche: a point in the hyper-dimensional plane of traits present in a given community. This concept of niche hyperspace concept was developed by Hutchinson (1961) himself. Filled circles represent species, according to the colour legend on the right hand side. Black diamonds represent the normalised nutrient supply rates 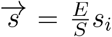. The green triangle represents the plane *α*_1_ + *α*_2_ + *α*_3_ = *ϵ*, which - as *E* = 1 - intersects the maximal uptake rate traits (*A*_1_, *A*_2_, *A*_3_) for each resource in the points *A*_1_ = (1, 0, 0), *A*_2_ = (0, 1, 0) and *A*_3_ = (0, 0, 1). Because of the assumption of metabolic trade offs, each species intersect the niche surface.

#### Model

The growth of each species is given by the growth function *g_σ_*, which describes the growth rate of each species as a function of nutrient uptake, *r*_*i*_(*c*_*i*_). It is given by the sum of the rates of each resource, *c*_*i*_, consumed by species *σ*.

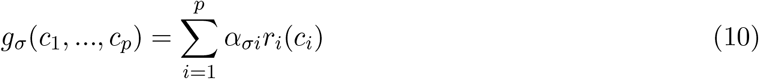

### 2.2 Modelling Microbial Communities - Demes

The mesoscale level of the model ecosystem comprises communities of species growing on homogeneouslysupplied, shared limiting resources.

#### Definition

A microbial community is constructed as an instance of the `Deme` class, defined by the concentration of shared resources 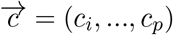, their rates of supply, 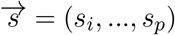 and the species that compose it, 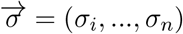.

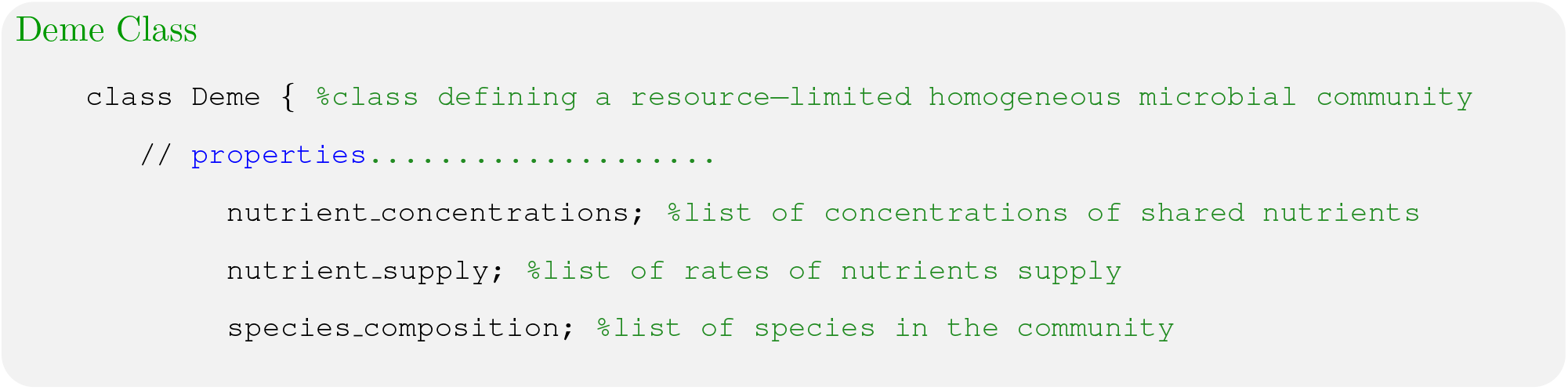

#### Assumptions

I. **Spatial Homogeneity:** Each deme is assumed to be a well-mixed system such that the concentration of nutrients is homogeneous and is determined by the nutrient supply rates 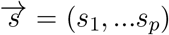, by the uptake of nutrients by organisms *r*_*i*_, and by a degradation or loss rate *μ*_*i*_.
II. **Resource-Competition Dynamics:** The dynamics of species and nutrients is described by a resource-competition model that incorporates classical generalised Lotka-Volterra ODEs and metabolic trade-offs, as recently described by (Posfai et al. 2017).
III. **Separation of Time Scales:** Since metabolic reactions typically occur on a faster time scale than cell division, these time scales can be separated, resulting in nutrient concentrations satisfying the flux-balance equation 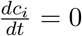

#### Model

In each deme, the population dynamics for each species is given by a set of ordinary differential equations (ODEs) describing each species growth, where the original “increase factor” *ϵ*, described by Volterra (1928) is replaced by the growth function for each resource consumed, *g*_*σ*_(*c*_1_, …, *c*_*p*_).

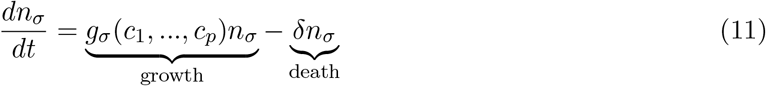

The kinetics of nutrient (*c*_*i*_) concentration is described by:

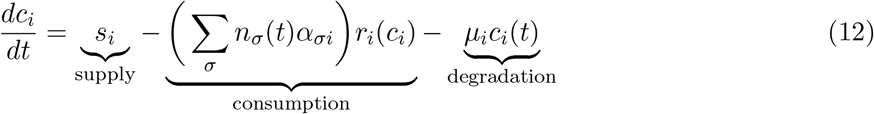

### 2.3 Modelling Structured Microbial Metacommunities

At the macro-scale, the ecosystem can be thought of as a “community of communities” (Wilson 1992), or metacommunity. The ecosystem can then be modelled as a matrix of communities (demes). Since phytoplankton lives floating on water, and since the fluid dynamics in the ocean is turbulent, species and resources can diffuse to other communities in the grid. This concept of metacommunity, is convenient as it allows to take the patchy structure of real ecosystems into account, without requiring computationally-intensive multidimensional simulations using partial differential equations (PDEs).

#### Definition

The ecosystem is modelled as a 2-dimensional discrete lattice of demes. It is defined as an instance of the `Plankton` class, and is constructed by specifying the demes that compose it and the dimensions of the lattice.

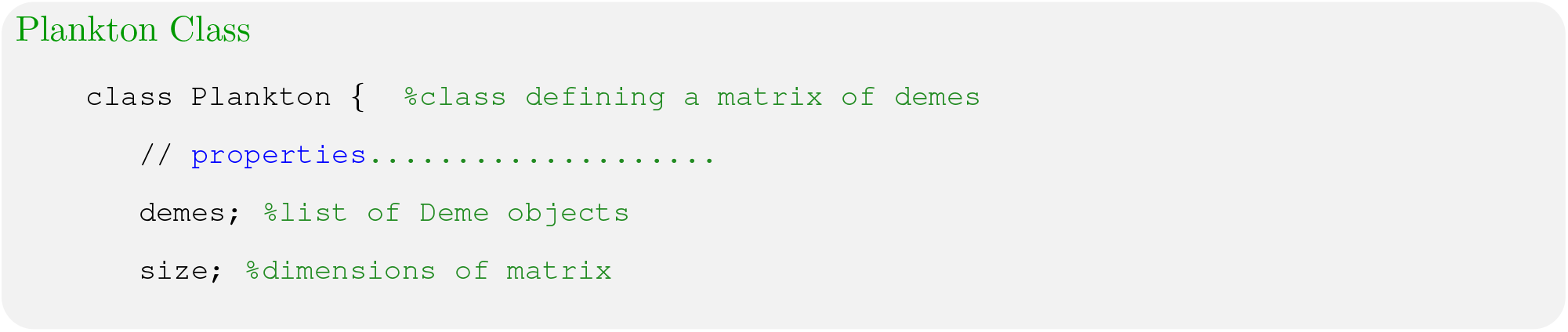

#### Assumptions

I. **Spatial Heterogeneity:** The distribution of resources is assumed to be heterogeneous (as shown in Figure 2). Every deme has the same total supply of resources, *S* = *s*_1_ + *s*_2_ + *s*_3_ = 1
II. **Random Dispersal:** Since communities are open systems, species can diffuse to other demes in the lattice. The probability of a species diffusing from one community to another is assumed to be inversely proportional to the Euclidean distance between the two communities in the lattice.
III. **Neutrality:** All species in the metacommunity obey to the same rules of ecological engagement, such as mixing and dispersal.

#### Model

The dynamics of the metacommunity is modelled by a discrete map of the species in each deme (Geyrhofer & Brenner 2018). As illustrated in Figure 4, every deme within the community is simulated independently according to Equation 11 and 12 until a steady state is reached, at a time *t*. Species are then allowed to diffuse to other demes in the lattice. The composition of each deme is then updated with the new invader species. Demes are simulated again and this process iterated until the total simulation time, *T*.

**Figure 4:**
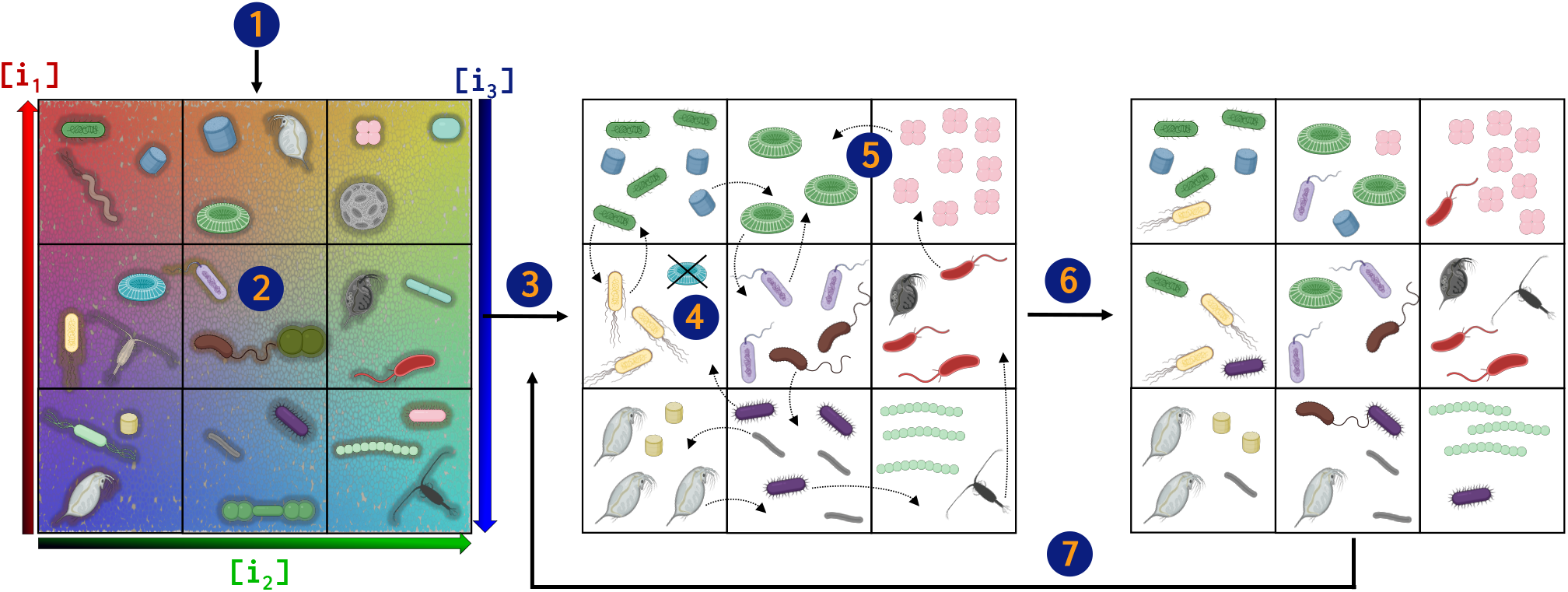
Schematic of the simulation algorithm. Numbered circles in the drawing represent the following steps: 1. The plankton ecosystem is initialised as a matrix of spatially isolated microbial communities (demes). Each deme contains *p* = 3 resources, with concentrations that vary from deme to deme to account for spatial heterogeneity, according to colours (*i*_1_ = red, *i*_2_ = green, *i*_3_ = blue). 2. Demes are initialised with *n* species. The traits of these species are randomly drawn from a uniform distribution, 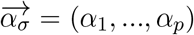. 3. Solve concentration and population dynamics ODEs in each deme according to Equation 16 and 17. 4. Species whose population value falls below a reference cutoff value (10^−5^) are removed from the global pool (black cross). 5. Species are allowed to diffuse across all demes. Diffusion is modelled as a random walk in a 2-dimensional array. The probability for a species to diffuse between two demes, donor and recipient, is inversely proportional to the distance, *d*, between the two demes, 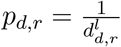. The distance between demes corresponds the Euclidean distance matrix representing the spacing between points in Euclidean space. 6. Every deme is updated with new species, thus new traits and new populations. 7. Iterate until iteration = T

## 3 Results

The biophysical framework described above was incorporated into the “plankton-simulator”, a set of MATLAB scripts that simulate the model according to the algorithm described in Figure 4 and in Appendix 7.2.6. The commented scripts, alongside the data presented below, are available on GitHub (https://github.com/scaralbi/plankton-simulator). Supplementary data and a guide on how to use the simulator can be found in the Appendix.

### 3.1 Reference Simulation

In order to compare results with varying parameter values and different assumptions, this section shows the results of a reference simulation. The parameters used are (listed in Appendix 7.2.7) as described by Posfai et al. (2017), which is used as a benchmark for the model developed in this thesis.

#### Question

What are the population dynamics in a spatially heterogeneous ecosystem with *p* = 3 resources?

#### Hypothesis

A spatially heterogeneous and stochastically-assembled metacommunity results in *m* >> p coexisting species.

#### Results

As shown in Figure. 5, after 200 generations, 8 species coexist in the metacommunity. Since *p* = 3, Figure 5 demonstrates that coexistence of *m* >> p species is predicted by the model. Sudden extinctions events and the fluctuating trajectories highlight the stochastic dynamics of the model, which agrees with experimental plankton observations (Benincá et al. 2008).

**Figure 5:**
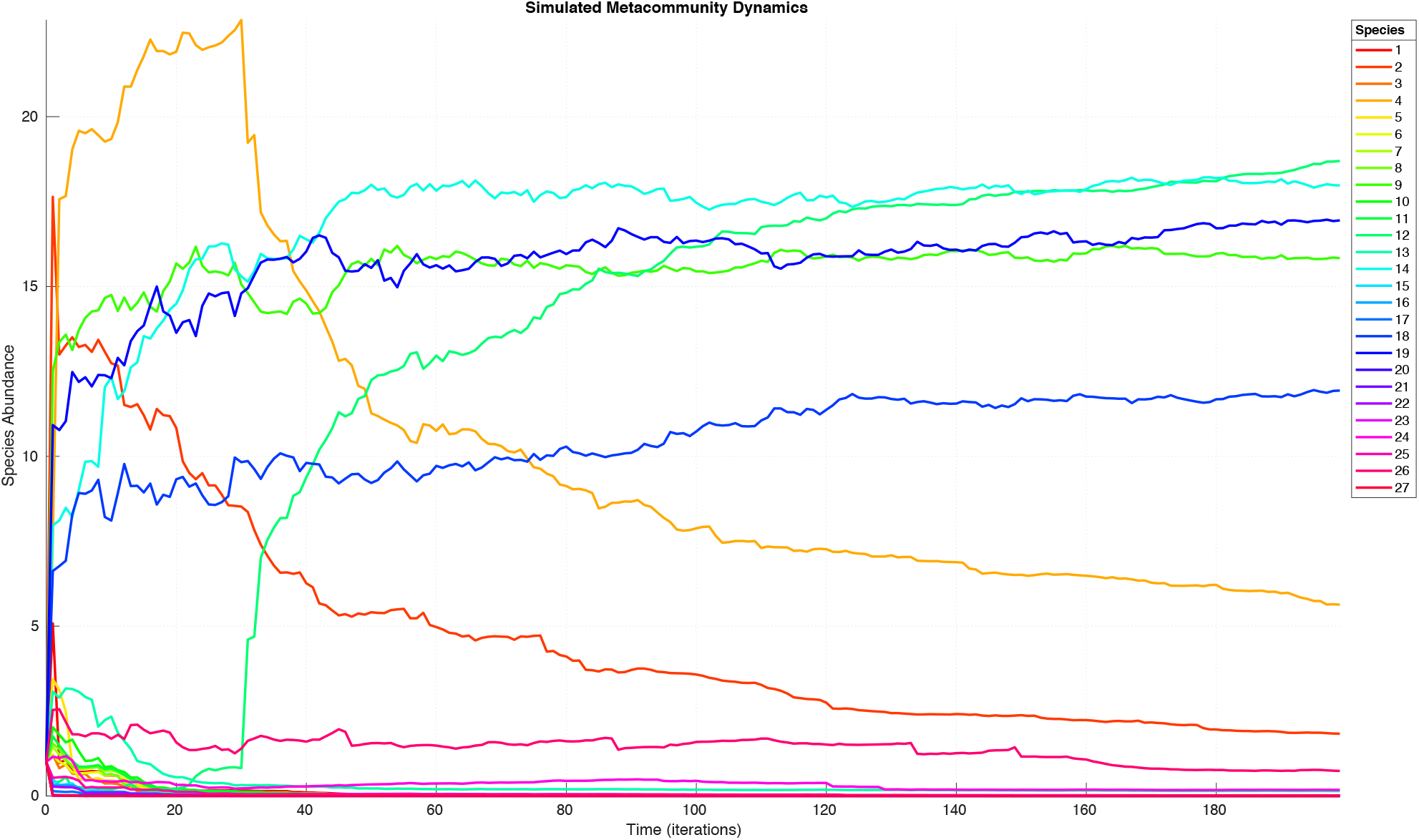
Population size of each species as a function of time (up to 200 generations). Species abundance is expressed in arbitrary units. Time is expressed in number of iterations. With the reference parameters used, every iteration corresponds to a generation time of *t* = 2000 seconds, thus resulting in *T* ≈ 4.6 days. Every species is assigned a colour according to the legend on the right hand side of the graph. A reference metacommunity is assembled as a 3×3 matrix of 9 communities. Every deme is initialised with 3 species drawn from a random pool of *m*_0_ = 27 traits 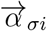. Each deme is supplied with *p* = 3 resources at rates 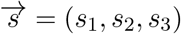. In every deme species grow until steady-state time, *t* = 2000 s. The simulation was run as described in the methods section for *T* = 200 iterations.

As it is assumed that all resources contribute to biomass accumulation, and energy is conserved, the total population size at steady state, 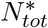, should satisfying the steady-state solution 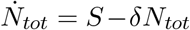, hence 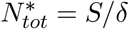. Indeed, since every deme is supplied with 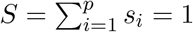 and there are 9 demes, it follows that the total supply in the metacommunity, *S*_*tot*_ = 9. Dividing *S*_*tot*_ by the reference death rate, *δ* = 0.01, results in 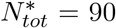. Interestingly, at *T* = 200, as shown in Table 2, the sum of the population values of the species present is 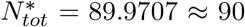. The reason for this approximation could be due to the use of the extinction cutoff described in the Methods section. Species with a population size below this cutoff are removed from the pool and not converted into resources (as opposed to what would happen in real ecosystems by the action of decomposer organisms). As a result, biomass is lost in the metacommunity, potentially explaining why the simulated 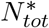 is lower than the value expected theoretically.

**Table 1.** Abbreviations and Symbols

| Table 1. Abbreviations and Symbols |  |  |
| --- | --- | --- |
| Abbreviation | Meaning | Reference Value |
| $\sigma$ | type of species | $\sigma_1, \dots, \sigma_{27}$ |
| $i$ | type of nutrient | $i_1, \dots, i_p$ |
| $p$ | total number of resources | 3 |
| $m_0$ | number of species initialised | 27 |
| $m$ | number of coexisting species | N.A. |
| $n_\sigma$ | population size | N.A. |
| $n_{tot}$ | total species abundance in the community | 2 |
| $N_{tot}$ | total species abundance in the metacommunity | 2 |
| $d$ | number of demes | 9 |
| $s_i$ | rate of nutrient i supply | $[0, 1]$ |
| $S$ | total supply rate | 1 |
| $\delta_i$ | death rate coefficient | 0.1 |
| $\mu_\sigma$ | nutrient degradation coefficient | 0.01 |
| $K_i$ | resource half-saturation constant | 1 |
| $t$ | species generation time | 2000$sec$ |
| $T$ | number of generations (total simulation time) | 200 (iterations) |
| $l$ | shuffle locality parameter | 2 |
| $\alpha_{\sigma i}$ | rate of resource utilisation | $[0, 1]$ |
| $\epsilon$ | rate of species increase | N.A. |

**Table 2.** Reference Simulation - Metacommunity Dynamics.

| Table 2. Reference Simulation - Metacommunity Dynamics |  |  |  |
| --- | --- | --- | --- |
| $p$ | $m_0$ | $m$ | $N_{tot}^*$ |
| 3 | 27 | 8 | 89.9707 |

The dynamics of species and resources at the scale of local communities can be analysed by plotting the population size of each species in each deme as a function of time, as shown in Figure 6.

**Figure 6:**
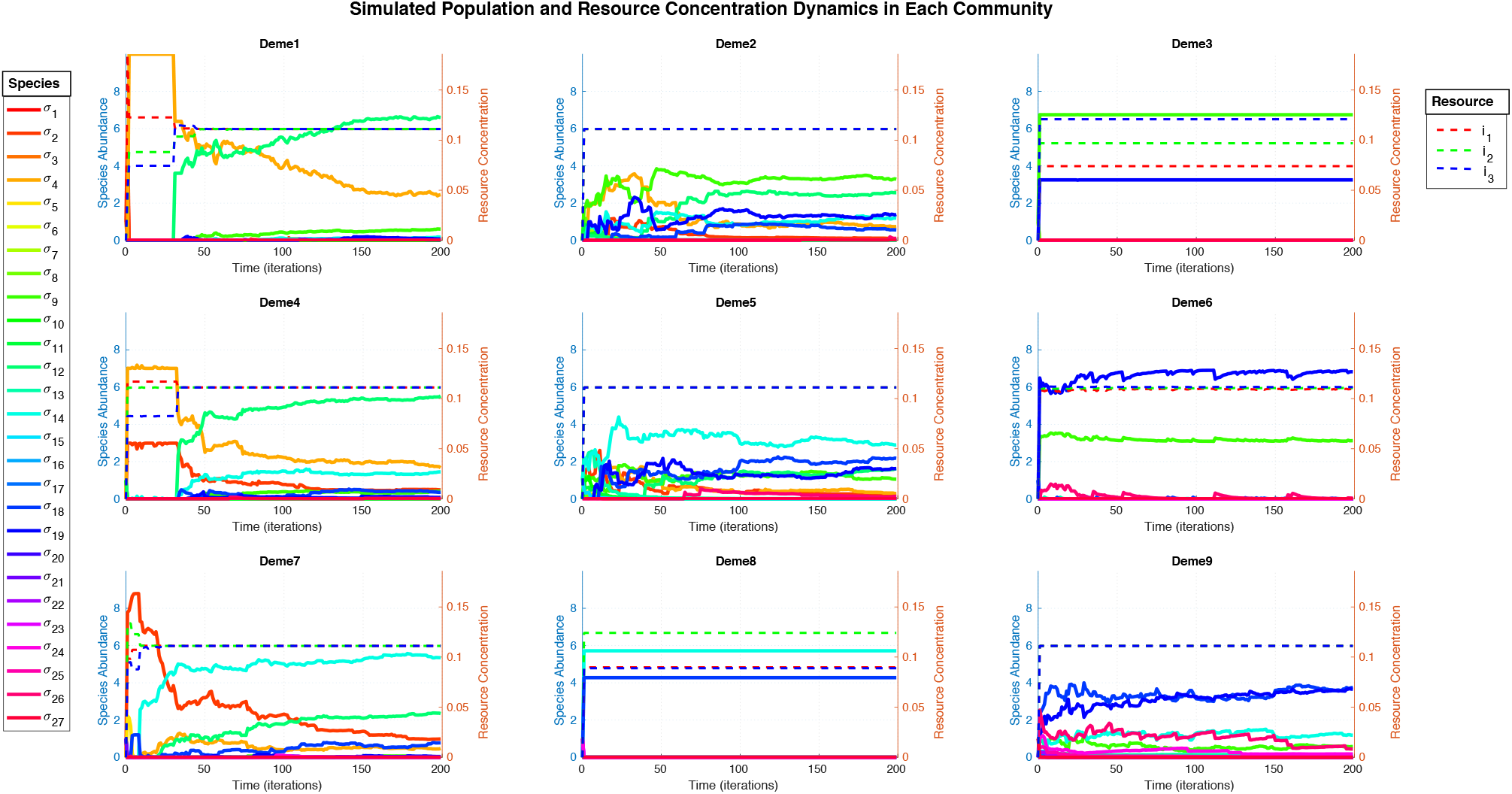
Species abundance and resource concentration as a function of time (up to 200 generations) in each community. Solid lines represent species according to the legend on the left hand side. Dashed lines represents resources, according to the legend on the right hand side. The left-hand side axis of each subplot illustrates the scale of species abundance (arbitrary units), while the right hand side the scale of resource concentration (arbitrary units). Every deme is supplied with a vector of resources, 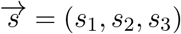 that varies from deme to deme as shown in Figure 2 to account for spatial heterogeneity.

The growth curves for each species have a similar hyperbolic shape as shown in Figure 5, asymptotically reaching the maximum population size determined by the carrying capacity of each deme. Following the same reasoning as above, the total cell abundance should satisfy the steady state solution, 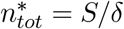. Indeed, Table 3 shows that the predicted 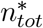 in each deme reasonably approximates to 1/0.1 = 10.

**Table 3.** Reference Simulation - Community Dynamics.

| **Table 3. Reference Simulation - Community Dynamics** |  |  |  |  |  |  |  |  |  |
| --- | --- | --- | --- | --- | --- | --- | --- | --- | --- |
|  | Deme 1 | Deme 2 | Deme 3 | Deme 4 | Deme 5 | Deme 6 | Deme 7 | Deme 8 | Deme 9 |
| $p$ | 3 | 3 | 3 | 3 | 3 | 3 | 3 | 3 | 3 |
| $m_0$ | 27 | 27 | 27 | 27 | 27 | 27 | 27 | 27 | 27 |
| $m$ | 7 | 17 | 2 | 9 | 18 | 6 | 9 | 2 | 6 |
| $n_{tot}^*$ | 9.967 | 9.9967 | 9.9971 | 9.9967 | 9.9967 | 9.9967 | 9.9967 | 9.9970 | 9.9967 |
| $c_1^*$ | 0.1111 | 0.1111 | 0.0741 | 0.1111 | 0.1111 | 0.1092 | 0.1111 | 0.0898 | 0.1111 |
| $c_2^*$ | 0.1111 | 0.1111 | 0.0969 | 0.1111 | 0.1111 | 0.1104 | 0.1111 | 0.1241 | 0.1111 |
| $c_3^*$ | 0.1111 | 0.1111 | 0.1209 | 0.1111 | 0.1111 | 0.1116 | 0.1111 | 0.0888 | 0.1111 |
| Convex Hull | Yes | Yes | No | Yes | Yes | No | Yes | No | Yes |

The resource concentration curves asymptotically reach a steady state (which can later be perturbed) after the first iteration. This is because the model has been simulated in discrete, integer, time steps (iterations) and as metabolic reactions happen significantly faster than cell growth, they satisfy the flux-balance assumption. Although every deme is supplied with different rates of resources at every iteration, deme 1, 2, 4, 5, 7 and 9 show that the concentration of resources reach a steady state such that 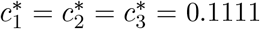. Interestingly, the aforementioned demes are the ones that are also populated by *m* > *p* species. This agrees with the findings of Posfai et al. (2017), who analytically demonstrated that if the normalised vector of nutrient supply, 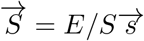, lies within the convex-hull (the pink dashed shape in Figure 7) of species traits, then the steady-state concentration of resources are 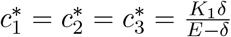. Substituting the reference parameters in this steady state condition results in 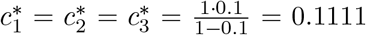, which corresponds to results shown in Figure 6 and Table 3. As illustrated in Figure 6, when this condition is met, coexistence of *m* > *p* is a robust outcome of the simulated dynamics.

**Figure 7:**
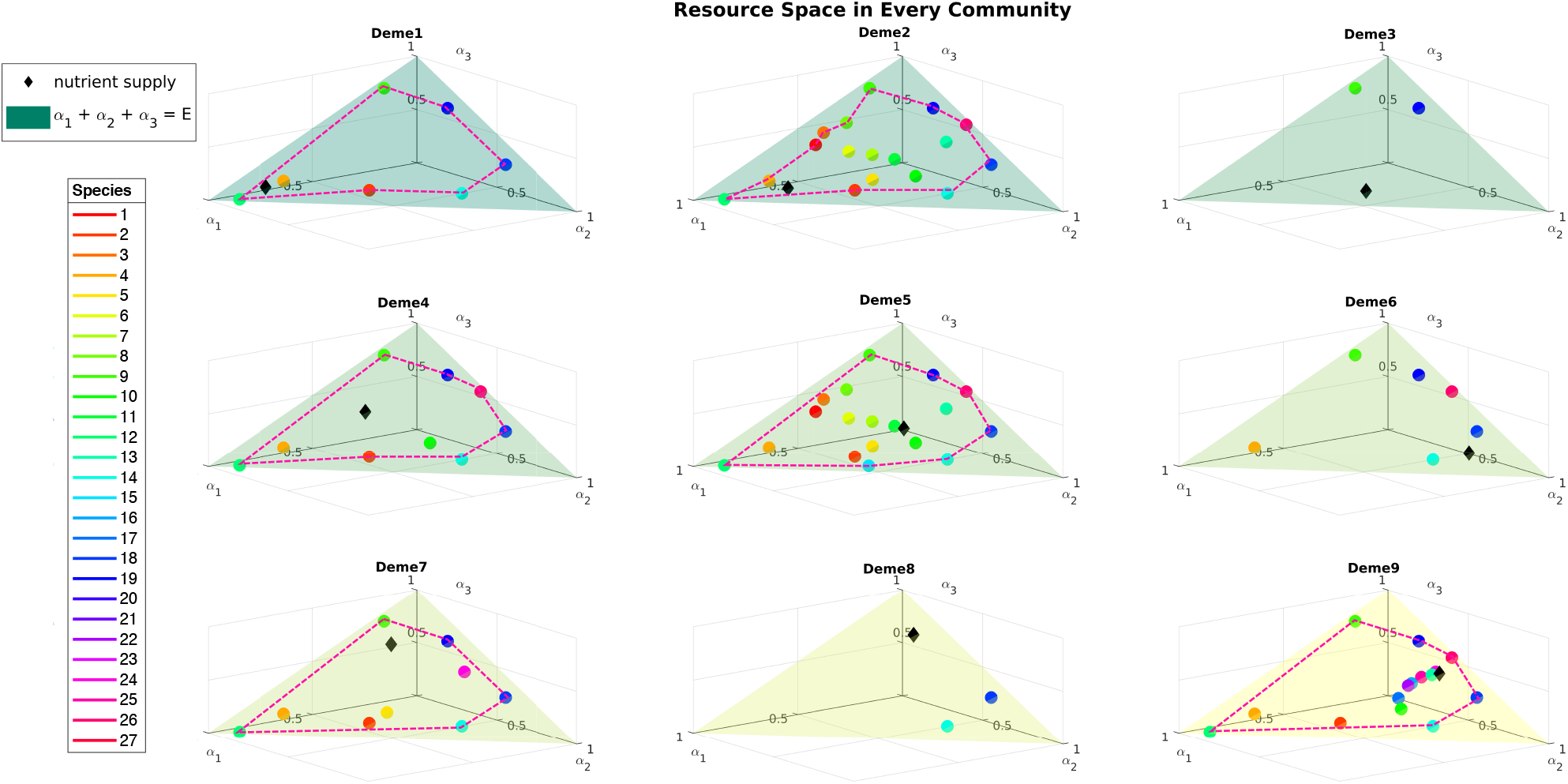
Since *p* = 3, every species can be plotted as a point in the 3-dimensional resource space defined by the metabolic traits 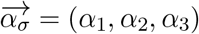. Filled circles represent species, according to the colour legend on the left hand side. Black diamonds represent the normalised nutrient supply rates, 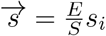. The green triangles represent the plane *α*_1_ + *α*_2_ + *α*_3_ = *E*, which - as *E* = 1 - intersects the maximal uptake rate traits (*A*_1_, *A*_2_, *A*_3_) for each resource in the points *A*_1_ = (1, 0, 0), *A*_2_ = (0, 1, 0) and *A*_3_ = (0, 0, 1). The pink dashed shapes represent the convex hull, that is smallest set of points on the niche surface that encloses the points occupied by each species.

By comparing the dynamics in Figure 6 and the distribution of species in the resource space (Figure 7 it can be observed in deme 1, 2, 4, 5, 7 and 9 that when the condition hull condition is satisfied 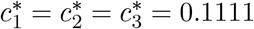, coexistence of *m* > *p* species emerges.

The population dynamics observed in Figure 5 and Figure 6 show that some species can suddenly go extinct or drive others to extinction. This is consistent with the simulation algorithm implemented, which at every iteration randomly draws some fraction of a species population (weighted by its current *n*_*σ*_) from a donor deme and updates it into a random recipient deme (inversely weighted by the Euclidean distance between the two demes). When a species invades a deme where another species close in the niche hyperspace is present, the more efficient one is expected to drive the latter to extinction. This indeed is illustrated in Figure 8, which shows the history of invasions across all demes in the metacommunity.

**Figure 8:**
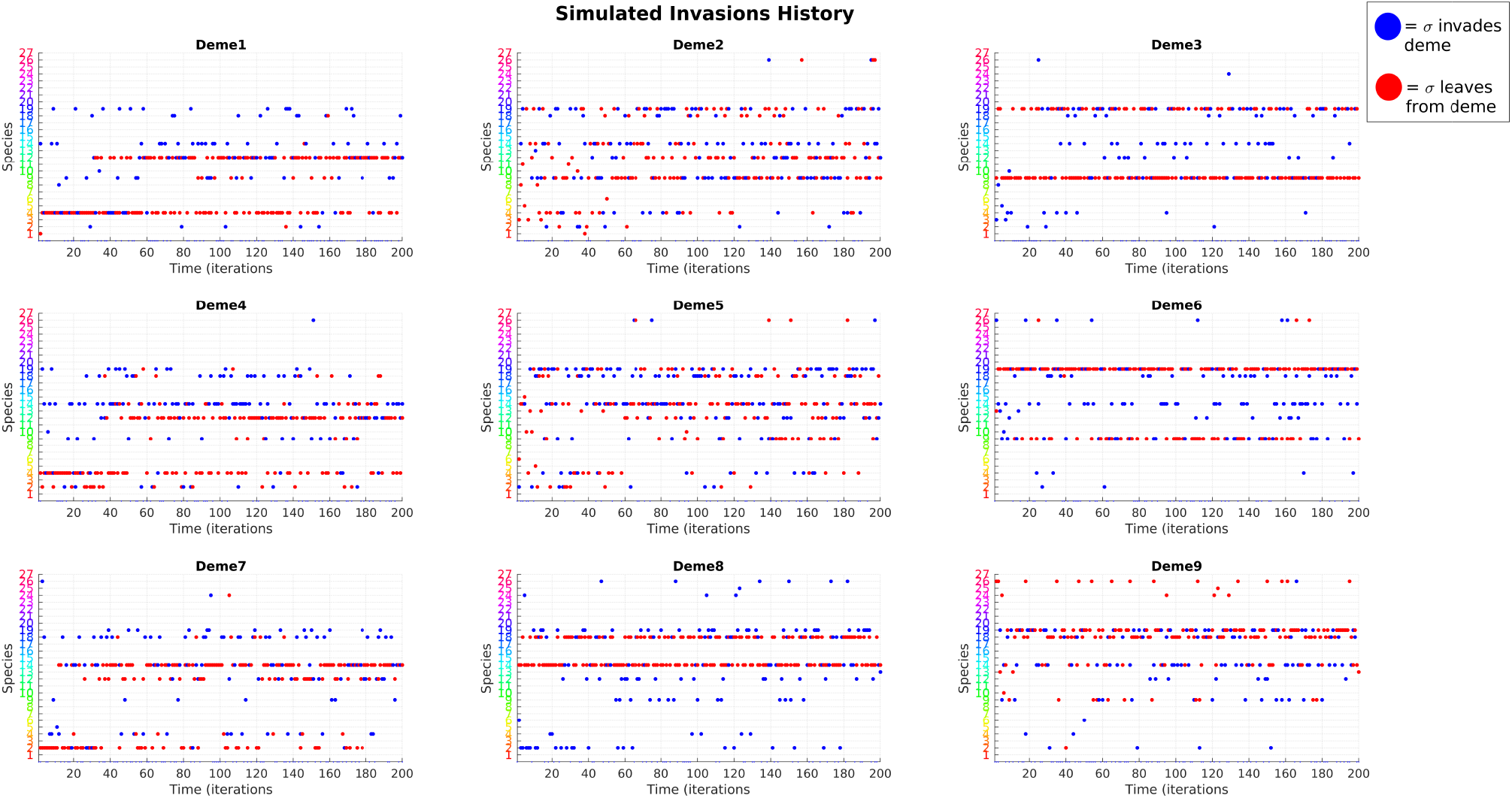
Every subplot corresponds to invasion events happening in a specific deme, as specified in each title. On the y-axis are shown the species, in ascending order by their names. The x-axis represents time in number of iterations. At every time step, a red point signifies that the species at the equivalent ordinate has been drawn as an invader in that deme. A blue point means that the species has invaded that deme at the time specified by the abscissa. Because invader species are randomly drawn weighted by their population value, a blue dot followed by repeated red dots (e.g *σ*_12_ in Deme 1) highlights that a species successfully invaded the deme, outcompeting the other species. Strikes of blue dots (e.g. *σ*_19_ in deme 1,4,7,8) suggest that the corresponding species is abundant in some other deme but is not particularly fit for the recipient deme. Species characterised by a similar ratio of blue and red dots (*σ*_14_) suggest that they are generalist species, as they can invade a deme, grow sufficiently to be drawn as invaders and then invade the deme again.

The history of invasions explains the sudden extinction and out-competition events in the simulated ecosystem. As an example, in Figure 5 it can be observed that the population of *σ*_4_ (orange line) dramatically falls at *T* = 30, whilst the population of *σ*_12_ (green line) suddenly increases. In fact, Figure 8 reveals that at *T* = 30, *σ*_12_ invades deme 1, while *σ*_4_ was thriving. As shown in Figure 7, deme 1 is mainly supplied by nutrient *i*_1_. Because *σ*_4_ is more efficient at uptaking *i*_1_ (as it is closer to the vertex *A*_1_ = (1, 0, 0), *σ*_12_ cannot compete and thus starts its decline to extinction. Other interesting cases can be analysed, whose descriptions however - for the sake of brevity - will be ignored here.

### 3.2 Effect of Perturbations in Metabolic Trade-Offs

The results above confirm that the simulation replicates the properties predicted by the (Posfai et al. 2017) model. The metabolic trade-off model explains how coexistence can emerge from the distribution of species traits in the resource space. However, even minimal perturbations of such traits result in dramatic loss of biodiversity, confirming the principle of competitive exclusion. In order to test if the model here proposed performs better than the aforementioned model, this section shows results of a modified version of the reference simulation, in which random variations to the metabolic traits are introduced.

#### Question

What are the effects of perturbing the metabolic trade-offs on the simulated population dynamics?

#### Hypothesis

A spatially heterogeneous and stochastically-assembled ecosystem is robust to perturbations in metabolic traits.

#### Results

As shown in Figure 9, similarly to the results from the reference simulation, after 200 generations, coexistence of *m* >> *p* species is predicted by the model.

**Figure 9:**
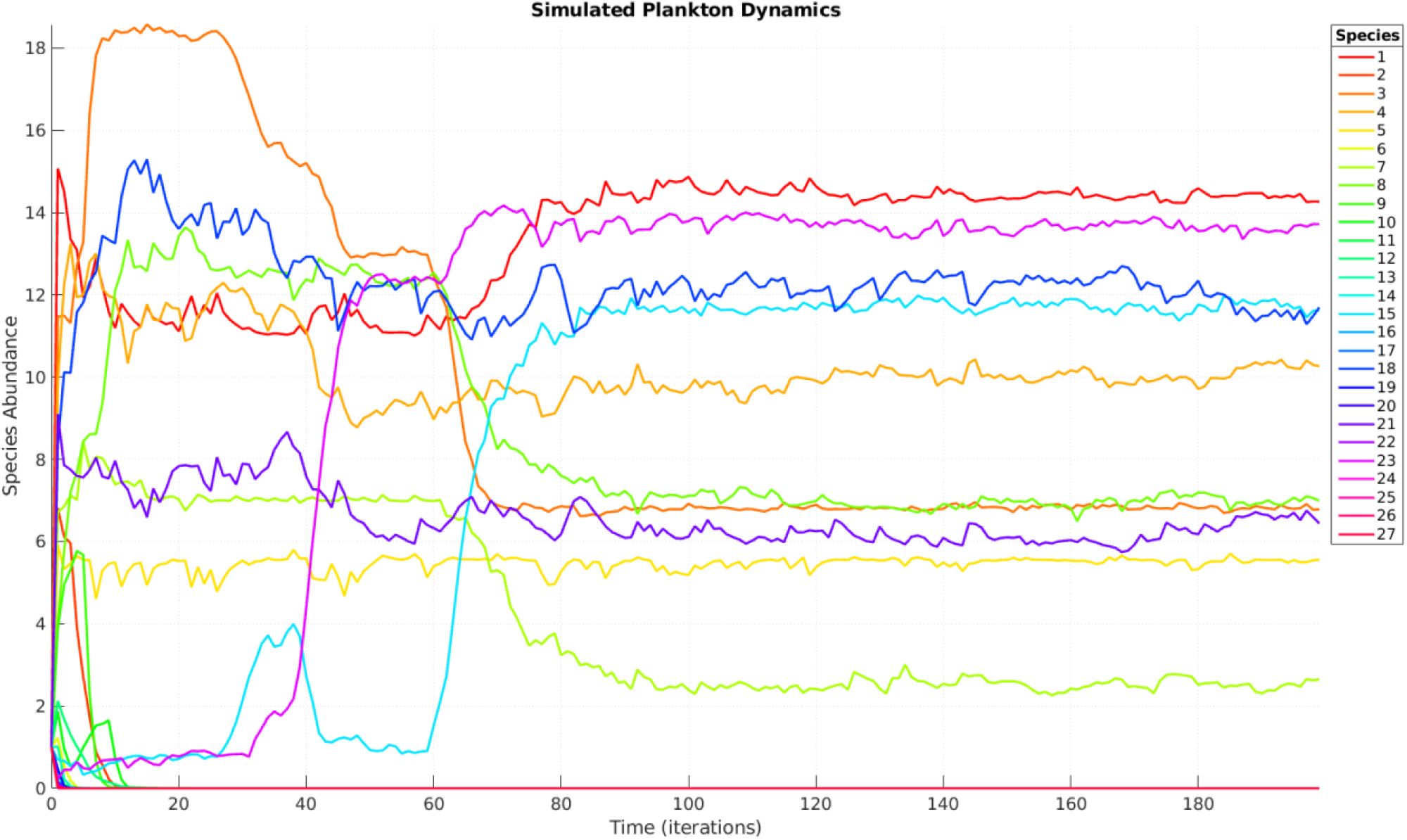
Population size of each species as a function of time (up to 200 generations). Species abundance is expressed in arbitrary units. Time is expressed in number of iterations. With the reference parameters used, every iteration corresponds to a generation time of *t* = 2000 seconds, thus resulting in *T* ≈ 4.6 days. Every species is assigned a colour according to the legend on the right hand side of the graph. To account for variations in *ϵ*, the vector of metabolic strategies is now 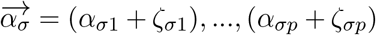 where 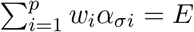 and *ζ_σi_* are independent random variables drawn from the Gaussian distribution 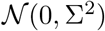, with standard deviation, Σ = 0.02, that is twice as large as the one reported by (Posfai et al. 2017). Perturbations are only allowed if *E* increases, otherwise species would inevitably go extinct. The simulation was run with the other parameters as described for the reference simulation.

The simulation is stochastic, thus results are different at every run. To test whether coexistence is a robust outcome of the model, the simulated ecosystem dynamics with perturbed trade-offs is compared with that of the reference simulation. Figure 10 illustrates the resulting patterns of species composition in the metacommunity, evaluated by plotting the relative abundance of species in descending rank order, in a so-called “rank-abundance” or Whittaker (1970) plot.

**Figure 10:**
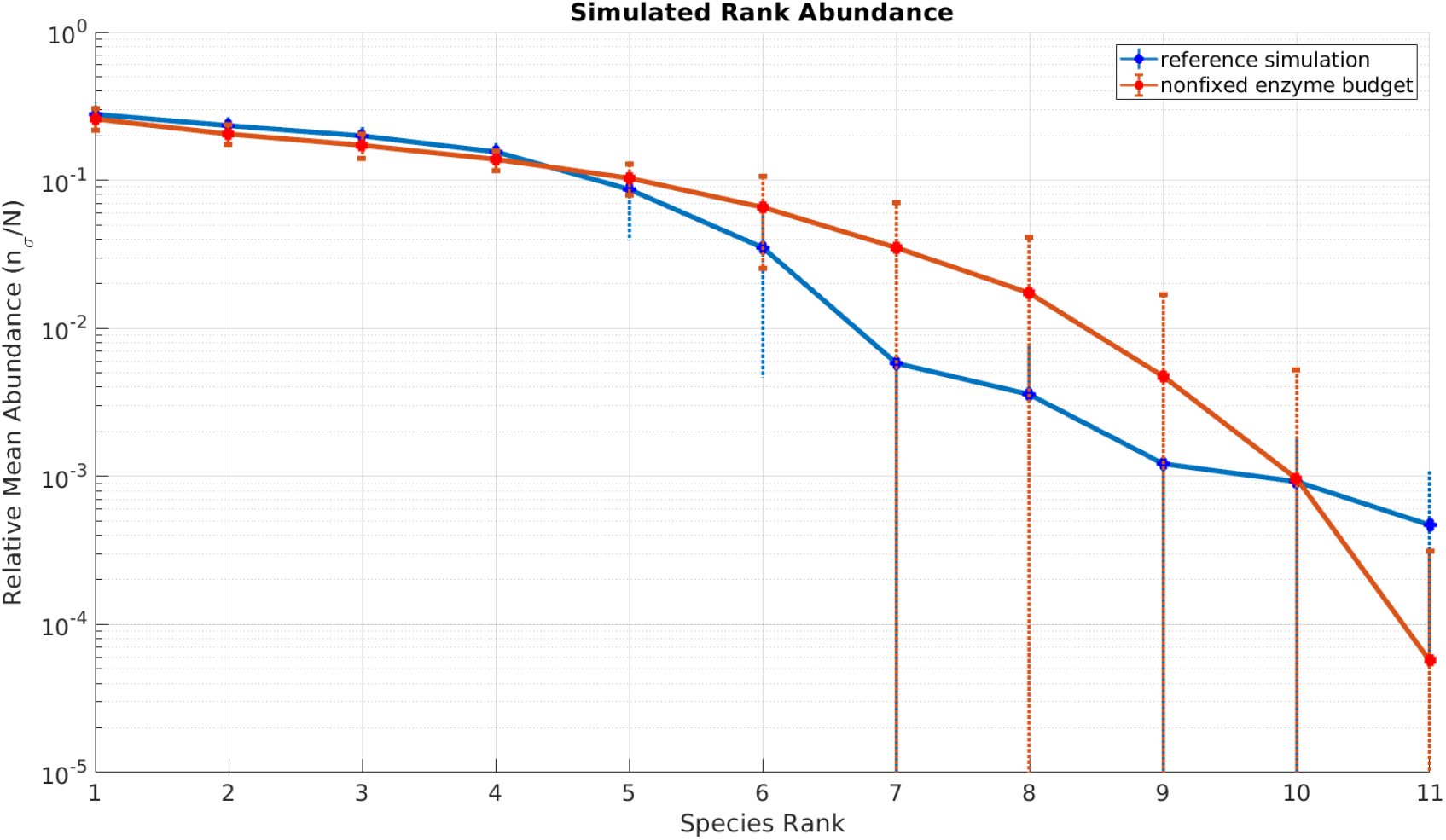
Rank abundances in the metacommunity at *T* = 200, simulated with reference parameters (solid blue line) and non-fixed enzyme budget. The y-axis is in *log*_10_ scale and shows the mean population size normalised by the total steady-state population, 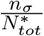 (unit-less). The x-axis corresponds to the species rank in the metacommunity in descending rank order (from the most common species to the rarest). The width of the plot corresponds to the number of coexisting species, while the slope shows the evenness of species distribution. The dotted vertical lines represent standard deviations 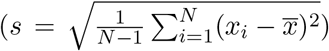 from *N* = 20 observations. From rank 7 symmetric error bars would lead to negative abundances, which cannot be plotted on the y-axis in log-scale. Therefore they are drawn until the x-axis, which intersects the y-axis at the cutoff value of 10^−5^.

The results show that the distribution of species abundances resulting from the reference simulation and the simulation with relaxed fixed enzyme budget assumption both approximate to lognormal curves, with mean value of 1 (the most common species occupies the larger fraction of the ecosystem). Until rank 6, there is no significant difference, confirming that even in the presence of perturbations in fixed-enzyme budget, the model predicts *m* > *p* species coexisting after 200 generations. The two curves start to diverge from rank 7, where the curve representing the non-fixed enzyme budget simulation shows a more shallow gradient, suggesting increased biodiversity. However, as shown by the standard deviations, the differences are not significant enough to assert that the simulation with relaxed fixed enzyme budget assumption results in increased coexistence.

### 3.3 Model Robustness to Parameter Changes

After having tested and improved the model, the robustness of the model was assessed by introducing variations into other parameters. This also provides opportunities to investigate the factors most affect the emergence of coexistence in the metacommunity, thus leading to the paradox.

#### 3.3.1 Number of Resources

##### Question

“What is the relationship between biodiversity number of resources in the metacommunity?”

##### Hypothesis

Biodiversity increases with the number of limiting resources

##### Results

As shown in Figure 11, biodiversity quantified by the Shannon entropy increases with the number of shared limiting resources in the metacommunity.

**Figure 11:**
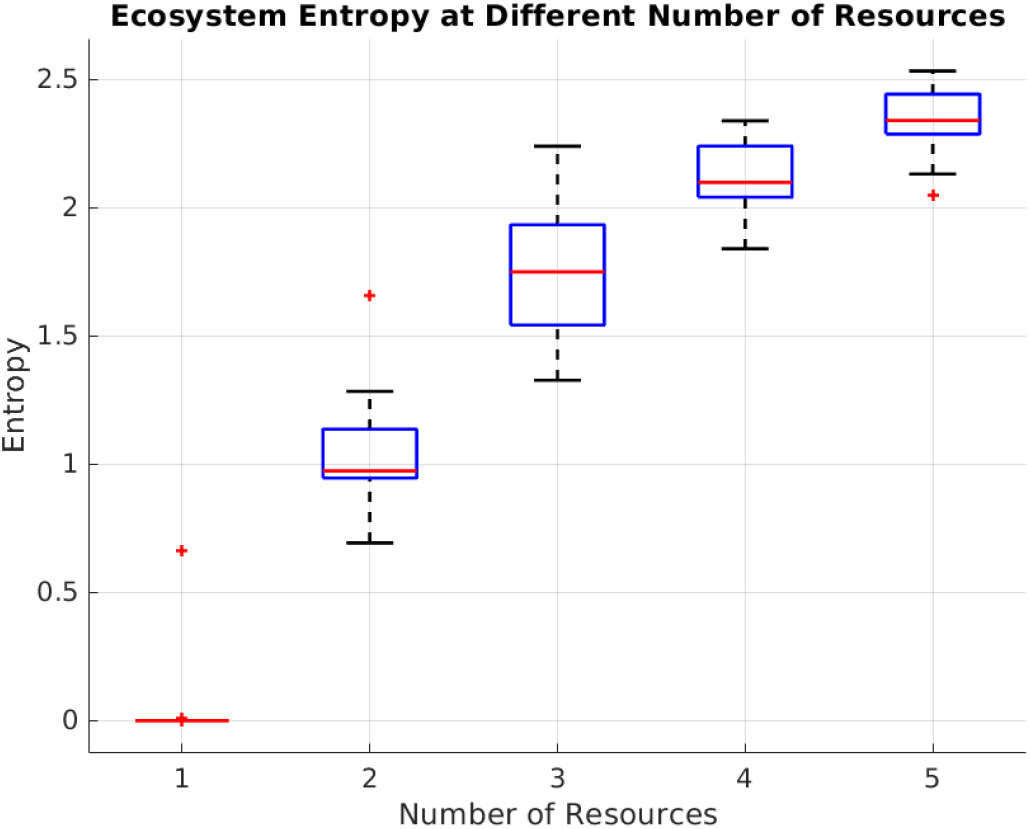
Shannon’s Entropy at different number of shared limiting resources. Every case was simulates 20 times according, as described for the non-fixed enzyme budget simulation. Entropy corresponds to the Shannon’s Entropy, 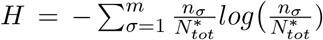 Units are in bits^−^1 ,On each box, the central red line represents the median value, and the top and bottom edges of the box show the 75th and 25th percentiles. The outliers are plotted individually with a + sign.

#### 3.3.2 Steady State Time

##### Question

“How does changing the mixing time, *t* affect the biodiversity of the metacommunity?”

##### Hypothesis

If species are shuffled across demes before they reach steady state, more biodiversity is expected.

##### Results

Figure 12 confirms the hypothesis by showing a two-fold increase in entropy when the the mixing time is changed from the reference value of *t* = 2000 sec to a value of *t* = 100 sec, which corresponds to the exponential phase of species growth in each community.

**Figure 12:**
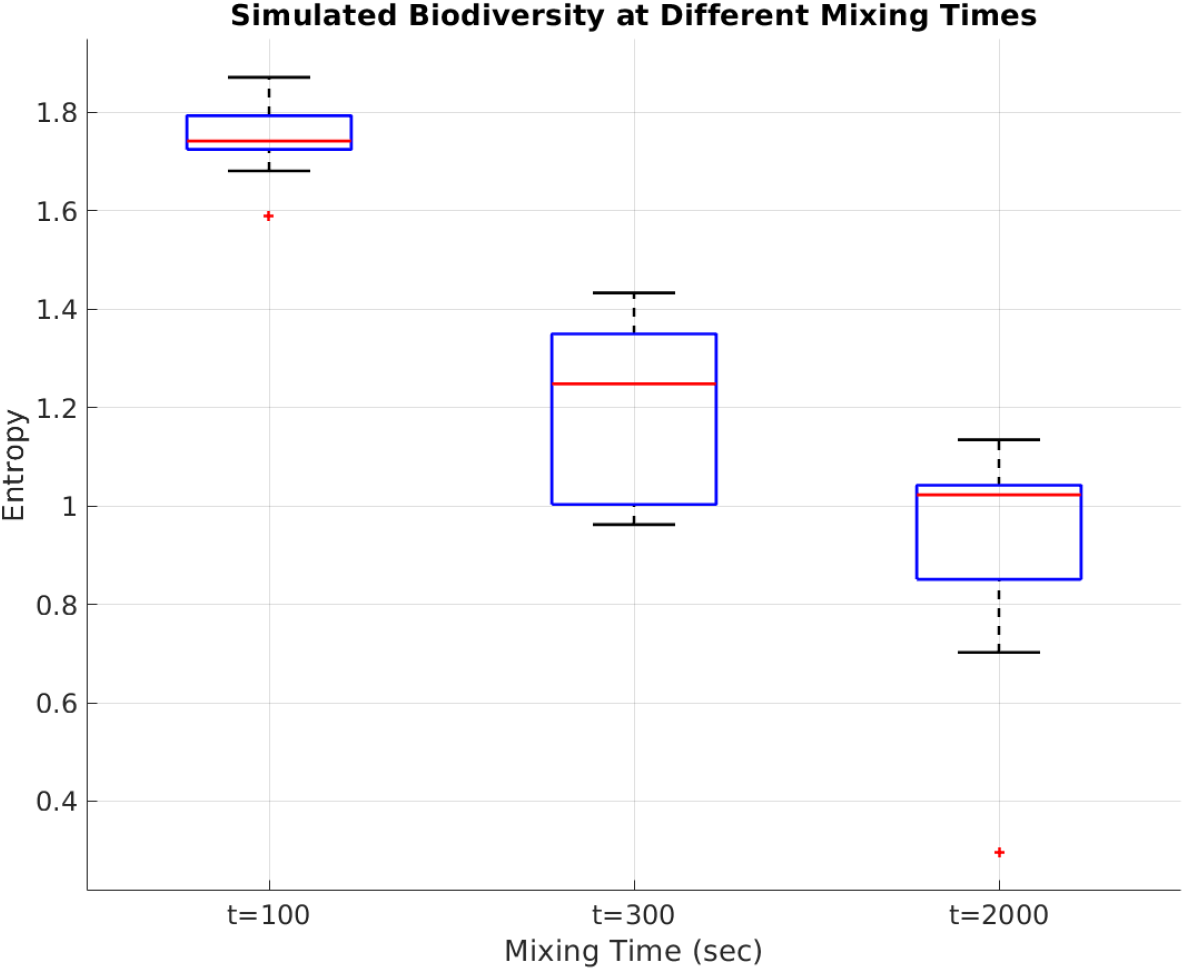
Metacommunity entropy at different mixing time, *t*. Biodiversity is quantified by the Shannon’s Entropy, 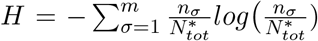 Every case was simulated 20 times according, as described for the non-fixed enzyme budget simulation. On each box, the central red line represents the median value, and the top and bottom edges of the box show the 75th and 25th percentiles. The outliers are plotted individually with a + sign.

Results show that the model predicts coexistence of *m* > *p* species at a wide range of mixing times *t*. This agrees with the similar model reported by (Huisman et al. 2001), which shows that when a community reaches a critical complexity to generate its own non-equilibrium dynamics, then *m* can exceed *p*. Therefore, the reason for the increased biodiversity at non steady state values in Figure 17 could be due to increased complexity in the ecosystem. However, although the model and the results are similar, the results are not enough to validate this hypothesis.

#### 3.3.3 Nutrient Supply

##### Question

“How do changes in nutrient supply affect the dynamics and stability of the ecosystem ?”

##### Hypothesis

Disruption of heterogeneity in the spatial distribution of resource is expected to result in loss of biodiversity and confirm the principle of competitive exclusion.

##### Results

As shown in Figure 13, significant changes are observed when the nutrient supply rates are set homogeneous.

**Figure 13:**
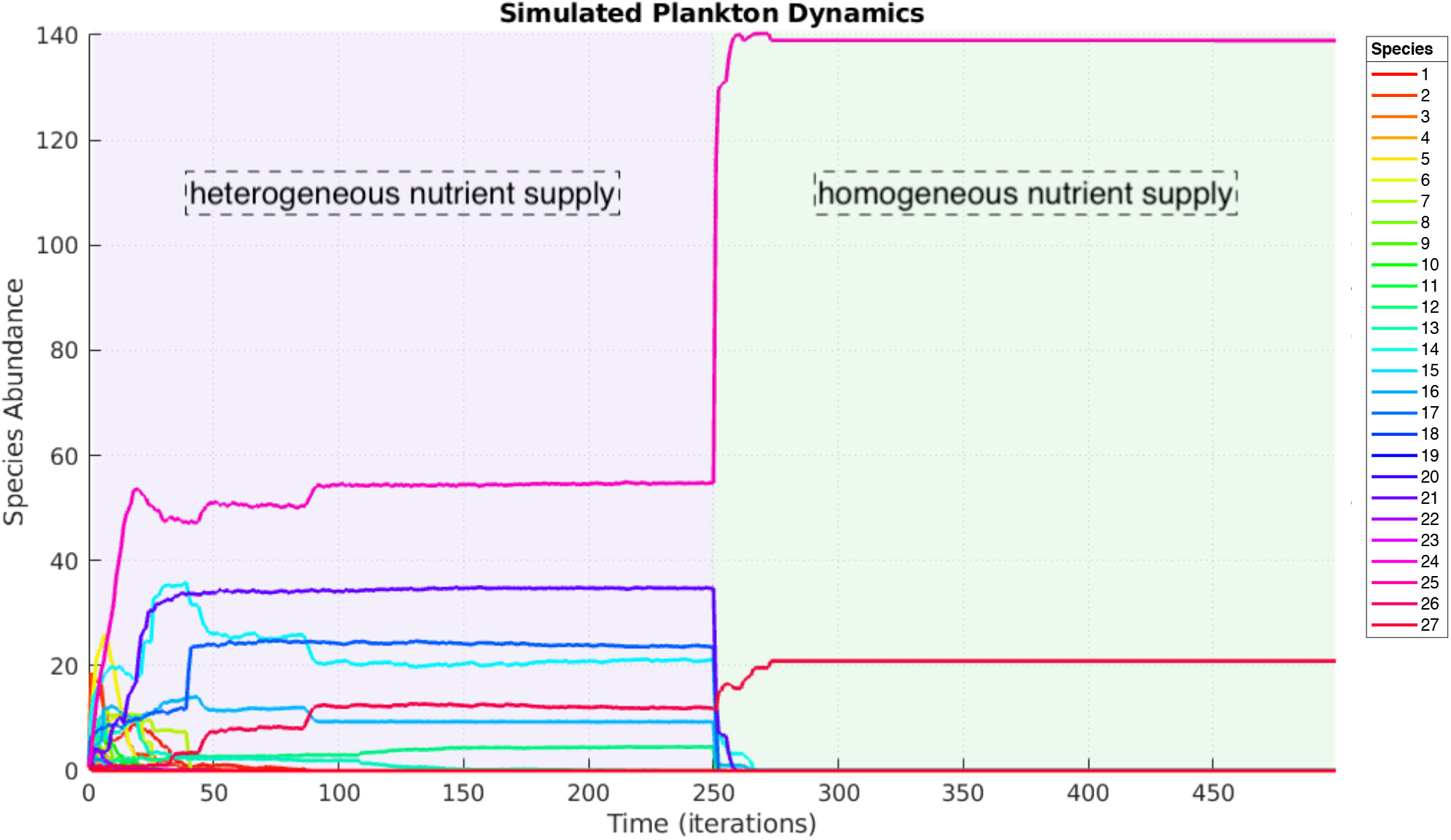
Population size of each species as a function of time (up to 500 generations). Species abundance is expressed in arbitrary units. Time is expressed in number of iterations. Every species is assigned a colour according to the legend on the right hand side of the graph.. For 0 < *T* < 250. the metacommunity is simulated as described for the non-fixed budget assumptions simulation. At *T* ≥ 250 nutrient supply rates are set to homogeneous values across all demes in the lattice 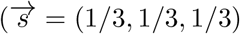 to simulate spatial homogeneity in the ecosystem.

When supply rates are set to homogeneous, species richness collapse from 8 surviving species to 2, thus confirming the principle of competitive exclusion. As shown in Figure 14, heterogeneity in nutrient supplies results in shallower gradients and reduced species richness compared to a homogeneously supplied ecosystem.

**Figure 14:**
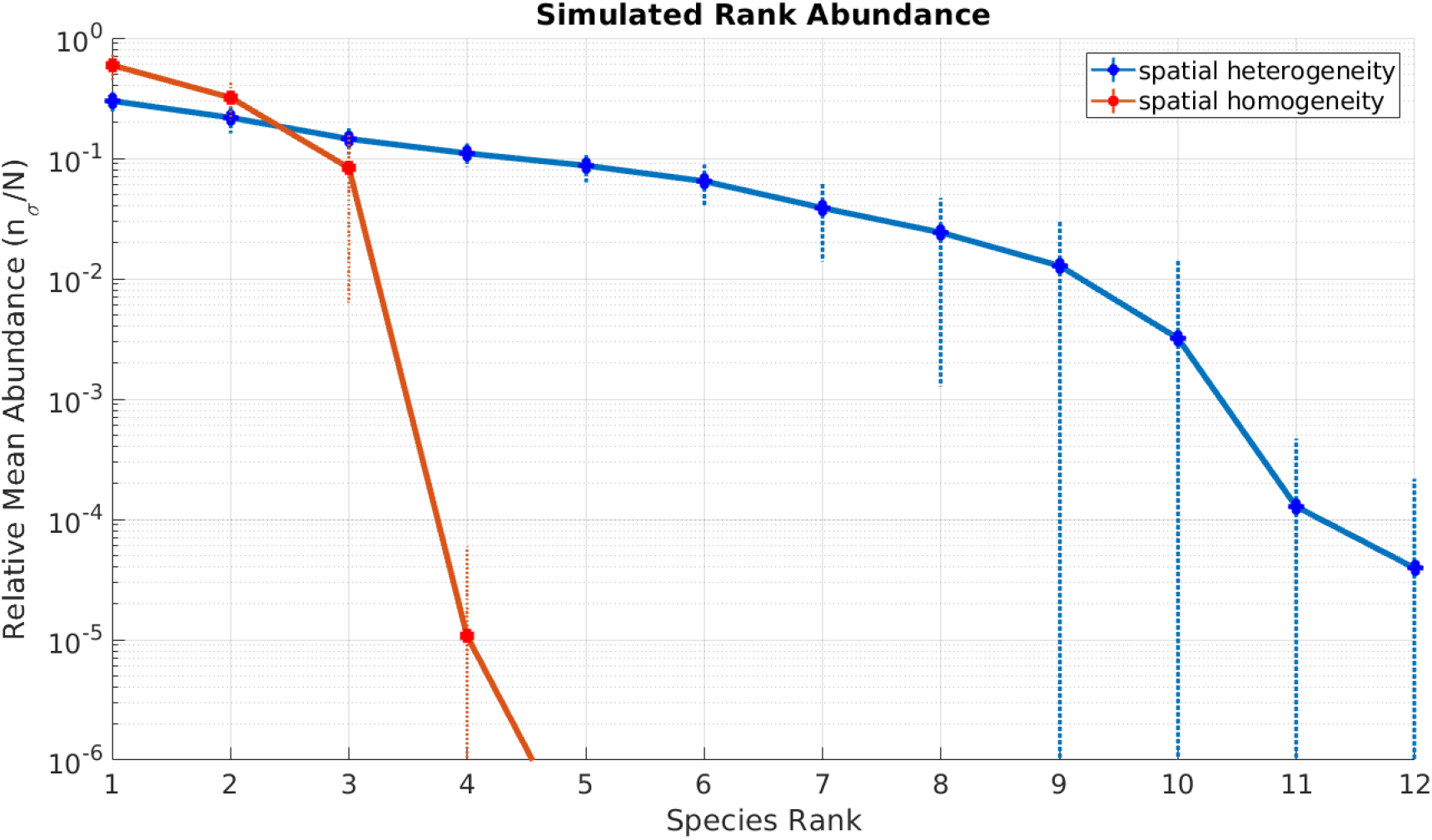
Rank abundances in the metacommunity at *T* = 200 (spatial heterogeneity, solid blue line) and at *T* = 400 (spatial homogeneity, solid red line), simulated without the fixed enzyme budget assumption. The y-axis is in *log*_10_ scale and shows the mean population size normalised by the total steady-state population, 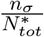 (unit-less). The x-axis corresponds to the species rank in the metacommunity in descending rank order (from the most common species to the rarest).The dotted vertical lines represent standard deviations 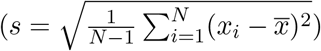 from *N* = 20 observations.

#### 3.3.4 Metacommunity Area

##### Question

“How does the area of the simulated metacommunity correlates with biodiversity ?”

##### Hypothesis

As Preston (1962) observed, species richness increases with increasing ecosystem surface.

##### Results

Species richness increases linearly as the number of simulated demes in the metacommunity increases.

At increasing number of demes, the slope of the rank-abundances curve become shallower, confirming the hypothesis of increased biodiversity. Indeed, Figure 16 shows that species richness increases linearly with the area.

**Figure 15:**
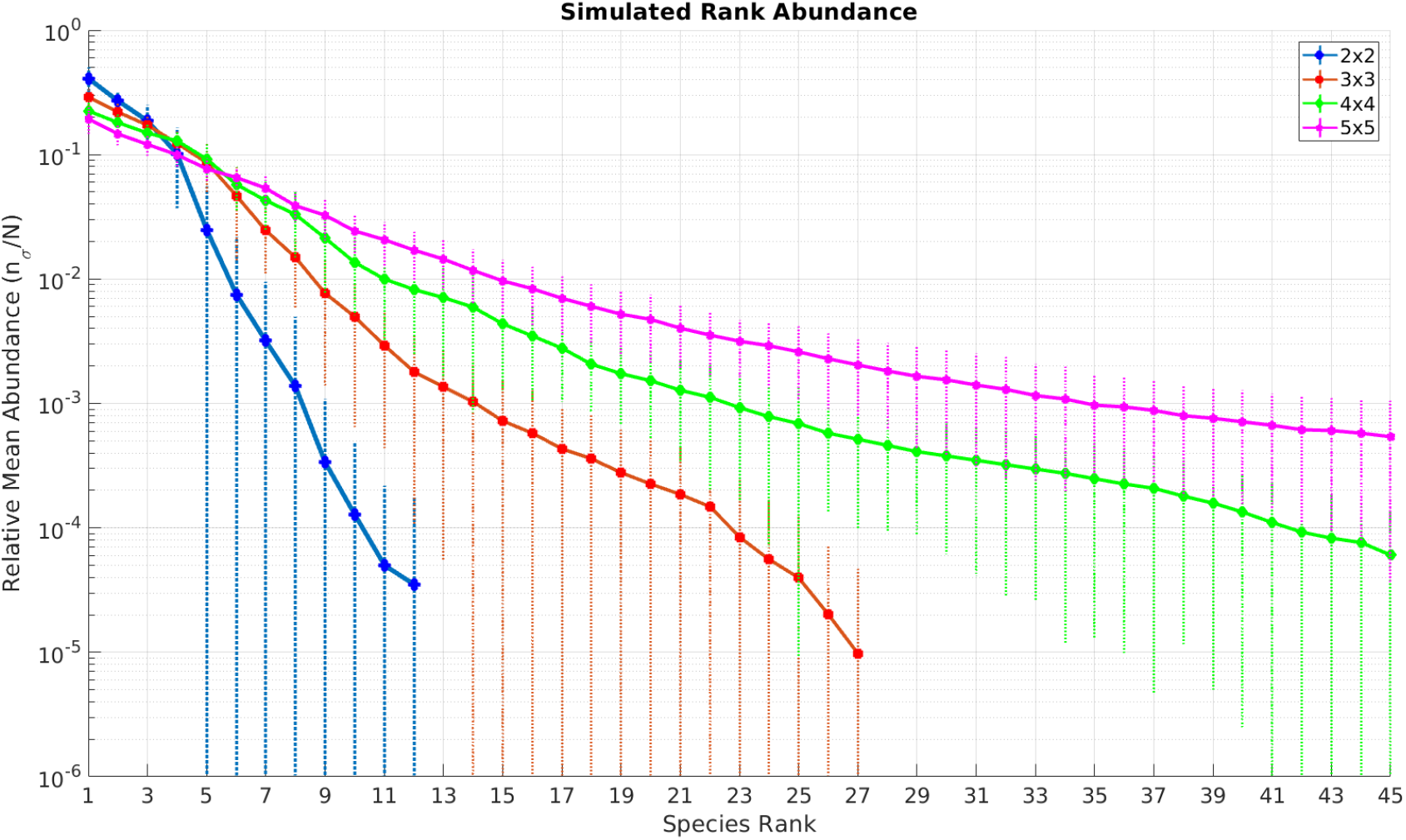
Rank abundances at increasing ecosystem area,at *T* = 200. The y-axis is in *log*_10_ scale and shows the mean population size normalised by the total steady-state population, 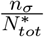 (unitless). The x-axis corresponds to the species rank in the metacommunity in descending rank order (from the most common species to the rarest).The dotted vertical lines represent standard deviations 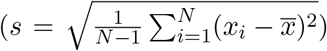 from *N* = 20 observations. In order to increase the surface area, simulations with non-fixed enzyme budget assumptions were run at increasing number of demes, to assemble metacommunities with dimensions specified on the legend on the right hand side (3×3 = 9 demes, 4×4 = 16 demes, 5×5 = 25 demes)

**Figure 16:**
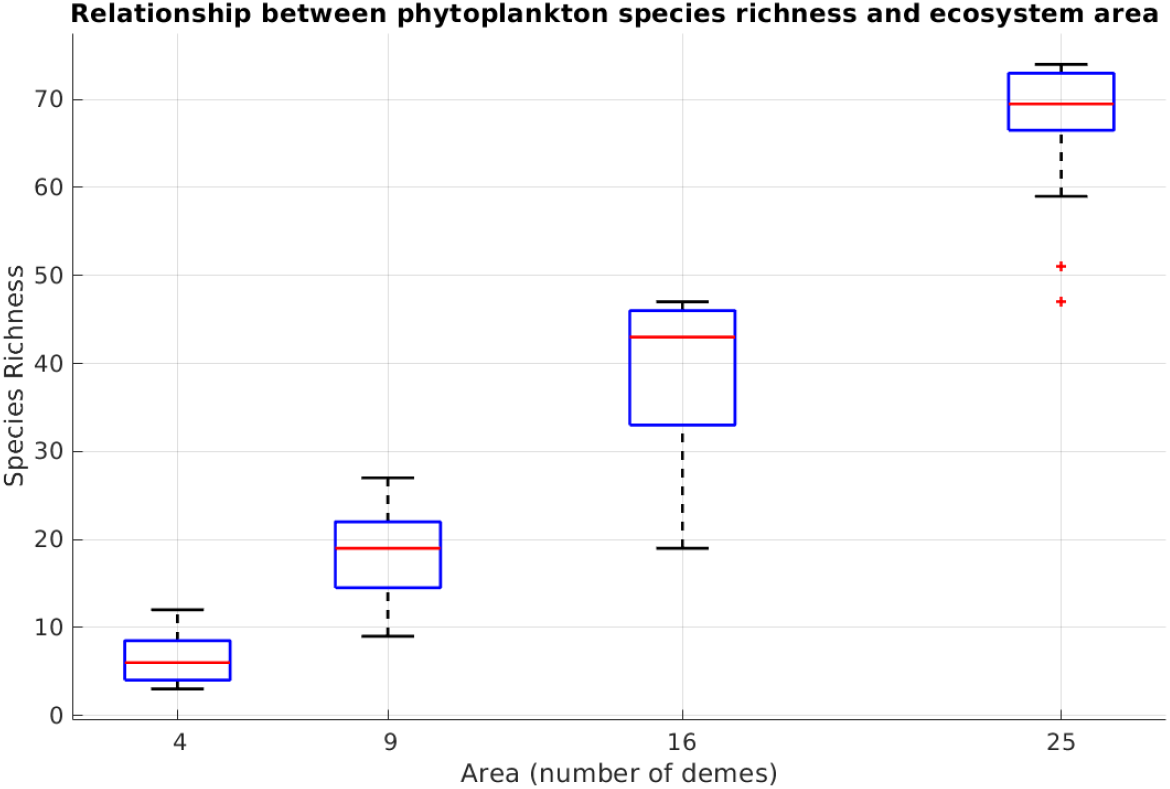
Species richness at different metacommunity areas. Species richness (y-axis) corresponds to the number of species coexisting in the metacommunity at *T* = 200 iterations. The metacommunity area (x-axis) is in number of demes. The simulation as described for Figure 15

The results agree with theoretical expectations from Preston (1962), who shows that species richness (S) increases linearly with the area (A) of the ecosystem. That is *S* = *j* · *A*, where j parameter which represents species density in each patch of community.

### 3.4 Model Validation

Finally, after having identified the parameters that most increase the simulated biodiversity, the model was compared with real data to investigate how the simulated pattern of species abundance compare with real phytoplankton distributions.

The simulated pattern of species has a similar lognormal shape as the distributions reported by (Margalef 1994). The simulated results predicts that the most common species in the metacommunity is less abundant than that reported by (Margalef 1994) and has a subsequent shallower slope. However, it has to be noted that, although widely used as comparison in the literature, the species in theis Margalef (1994) dataset were categorised from the microscope observations, thus potentially underestimating the real number of species present. Nonetheless, comparing the two curves confirms with reasonable approximation that species richness and average evenness of the simulated ecosystem match empirical phytoplankton observations.

## 4 Discussion

At the metacommunity level, results show that the proposed modelling framework predicts a larger number of species coexisting than that predicted by the principle of competitive exclusion. Population dynamics in the metacommunity (Figure 5) reveal that *m* > *p* species can coexist after ≈ 4.6 days in the model ecosystem. Since the simulation is stochastic, these results could be due to random immigration processes. This possibility is excluded by Figure 11 and 12, which predicts that total entropy of the metacommunity has a consistent median value of ≈ 1.2bits^−1^. In addition, Figure 17 confirms that the resulting lognormal distribution of species, consistent with the “canonical distribution of commonness and rarity” described by Preston (1962), fits experimental observations of real phytoplankton ecosystems made by Margalef (1994).

**Figure 17:**
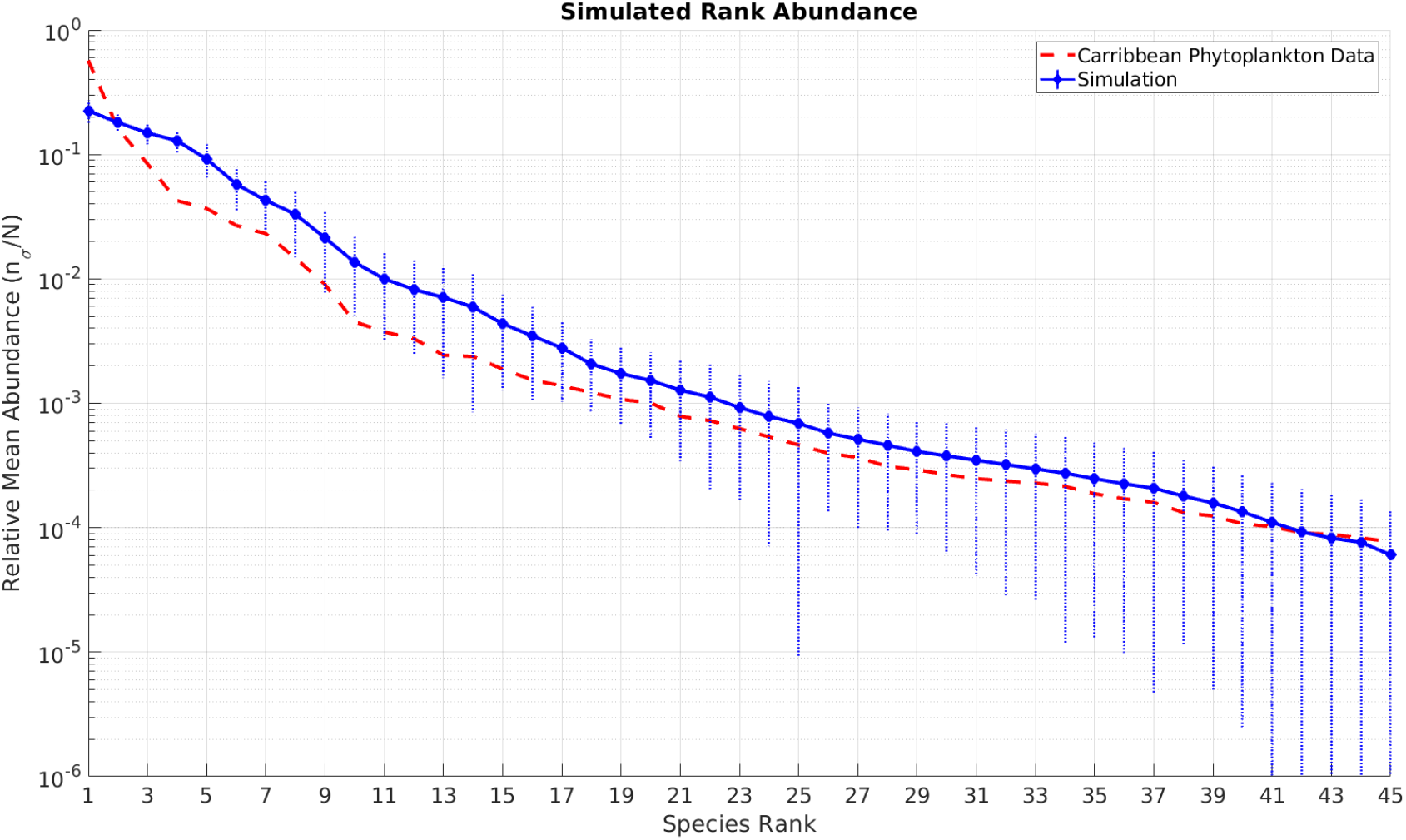
Rank abundances curves for simulation results (solid blue line) and real phytoplankton data (dashed red lines). The y-axis is in *log*_10_ scale and shows the mean population size normalised by the total steady-state population, 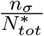 (unitless). The xacoxrisresponds to the species rank in the metacommunity in descending rank order (from the most common species to the rarest). The dotted vertical lines represent standard deviations 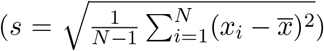 from observations. The real data are have been collected from the sea of Venezuela by (Margalef 1994), and are used as comparison also in (Laan & de Polavieja 2018). Each sampled volume was 150 ml, for a total surface of 20,000 km squared, and a total volume of 11.44 L. The simulation was run as described for the non-fixed enzyme budget assumption but with a 4×4 metacommunity of 16 demes, initialised with *m*_0_ = 45 species and *p* = 3 resources

In each local community, the number of species coexisting in each local community rarely exceeds the number of resources, apparently satisfying the principle of competitive exclusion. However, in the communities where the normalised nutrient supply, 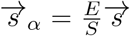, lies within the convex hull of species traits, 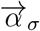, coexistence of *m* > *p* species is observed. This agrees with the findings of Posfai et al. (2017), who demonstrates analytically that when this condition is met, the hyperdimensional plane defined by species traits and nutrient supplies constitutes an attractor of the dynamical system. The model proposed here extends their findings by (1) considering a spatially heterogeneous ecosystem, and (2) showing robustness to variations in metabolic trade-offs. As species are subject to the same trade-offs in metabolic activities, a species that is very efficient at metabolising certain nutrient cannot be as efficient at metabolising another nutrient. Generalist species on the other hand are not efficient at metabolising any specific resource. However, since they are able to uptake a wide range of resources, they have an advantage when resources fluctuate and are able to inhabit a wide range of communities. Coexistence thus emerges as a result of the ability of species to sense, adapt and modify their surroundings.

Like in a welfare society, the distribution of wealth (enzyme budget) in the ecosystem is equally distributed among the population (because species are subject to same trade-offs). This allows species to “cooperate” in driving the amount of nutrients to concentrations for which species with different traits are equally fit (Laan & de Polavieja 2018). In addition, as expected from the assumption of neutrality in the ecosystem, the model also shows features of the neutral theory of biodiversity (Hubbell 2006). In fact, variations in the abundance of species are caused by stochastic processes of invasions and extinctions. In agreement with Litchman & Klausmeier (2008), this model predicts only patterns of diversity in the metacommunity, rather than taxonomic equilibrium. However, the role of niche differentiation remains the main factor determining the distribution of species in each community, thus potentially reconciling the two contrasting theories.

### 4.1 The Fallacy of the Plankton?

By comparing the distribution of species in spatially heterogeneous versus homogeneous environment Figure 14, the simulation clearly reveals that spatial heterogeneity significantly facilitates the emergence of coexistence. The findings agree with Dolson et al. (2017), who showed that spatial heterogeneity and biodiversity are positively correlated. The findings also agree with the simulations of competitive microbial communities performed by Lowery & Ursell (2019), who shows that the biodiversity in similar model ecosystem falls dramatically when environment with complex surface structures becomes isotropic. An explanation of these findings is proposed by the “positive heterogeneity-diversity hypothesis” (Stein et al. 2014), which states that spatial heterogeneity promotes diversity by creating new niches for specialist species to invade, thus limiting competition for shared nutrients. In other words, anisotropy in the ecosystem increases the dimensions of the niche hyperspace. The simulation also shows that even in stochastically-assembled, homogeneous environments, coexistence of *m* > *p* species is a robust outcome, if species collaborate to balance the concentration of resources in the environment. By maintaining classic Volterra (1928) resource-competition arguments but reevaluating some of its assumptions, the simple model here proposed matches empirical observations of real phytoplankton data and predicts robust coexistence of *m* > *p* species. The philosopher Quine (1976) categorises a paradox as a fallacy when a logical error is implied in the arguments. A paradox is categorised as falsidical when faulty reasoning is identified in the arguments. The results of this thesis suggest that, rather than a paradox, it is the consequence of false premises.

### 4.2 Future Perspectives

Models are only approximations of reality and, as such, this model relies on several simplifications. In particular, it focuses on the spatio-temporal dynamics of species solely defined by their metabolic traits. However, organisms possess an indefinite amount of them. For example, cell size here is not taken into account, which determines sinking rates and thus is an important factor in the spatial distribution of organism. Indeed, differential diffusivities between activator (preys) and inhibitor species (predators) might explain the emergence of spatial patterns of species distribution via Turing instabilities Tian & Zhang (2013). In addition, in our model every species is assumed to have the same generation time. In reality species have different replication rates, which might be important for oscillatory coexistence Guirey et al. (2010). Furthermore, species not only consume resources but also secrete organic matter as waste products, which can then be used as resources by other species. This cross-feeding mechanism is not taken into account here, and as reported by Goyal & Maslov (2018) might facilitate coexistence. Lastly, although the model simulates stochastic invasions and random evolution of traits, the long-term effect of evolution on the stability of the ecosystem is not incorporated. This could lead to significantly different results.

## 5 Conclusion

To summarise, a novel modelling framework was proposed, in order to simulate the dynamics of structured microbial metacommunities. By combining classic Lotka-Volterra dynamics (Volterra 1928, Hardin 1960) with trade-off consumer-resource dynamics Posfai et al. (2017) and stochastic ecosystem assembly (Geyrhofer & Brenner 2018), the model provided novel insights into the dynamics of structured microbial consortia. The simulation is modular and easily extendable, making it an ideal tool to predict changes in biodiversity due to anthropogenic climate change, identify and remove species that drive algal blooms (Hallegraeff 2010) and design synthetic microbial consortia for synthetic biology (Niehaus et al. 2019). In conclusion, this thesis demonstrated how a simple system-level simulation of a model ecosystem can be used as a heuristic tool to examine a theoretical framework - in this case that of competitive exclusion - and identify the fallacious assumptions that lead to its empirical falsification.

## 7 Appendix

## 7.1 Summary

The plankton-simulator is a collection of matlab scripts simulating the dynamics of structured microbial communities. The simulator is based on a multiscale biophysical model that incorporates species traits, consumer-resource dynamics and stochastic ecosystem assembly. The model hierarchically simulates features of ecosystem at different scales by defining three classes in the ecosystem using object-oriented programming in MATLAB: Species, Demes (spatially-isolated subcompartments) and Plankton (collection of Demes).

## 7.2 How to Use the Plankton Simulator

1. Download and extract the plankton-simulator GitHub repository; alternatively, clone the repository using Git with the command `git clone` https://github.com/scaralbi/plankton-simulator.git
2. Open MATLAB (this version has only been tested with version r2018)
3. Open `config.m`
4. Specify desired parameters (params), as described in file `params guide.tex`
5. Run `config.m`
6. Open and execute `simulator.m`
7. To visualise simulation plots and statistics execute `data reader.m`

## 7.2.1 Overview of the files in the plankton-simulator folder

- Species.m: defines the Species class and the functions that can be applied its objects
- Deme.m: defines the Deme class and its functions
- Plankton.m: defines the Plankton class and its functions
- dynamics.m: contains the ordinary differential equations that describe the population and concentration dynamics within ecosystem
- config.m: script that generate configuration files with speified parameters to run the simulation
- simulator.m: script that reads config files and simulate ecosystem and save data as config data.mat files
- data reader.m: script that reads config data.mat files and produces plots and statistics

## 7.2.2 Species.m

The Species class defines a single microbial species. Every Species is defined by the following properties:

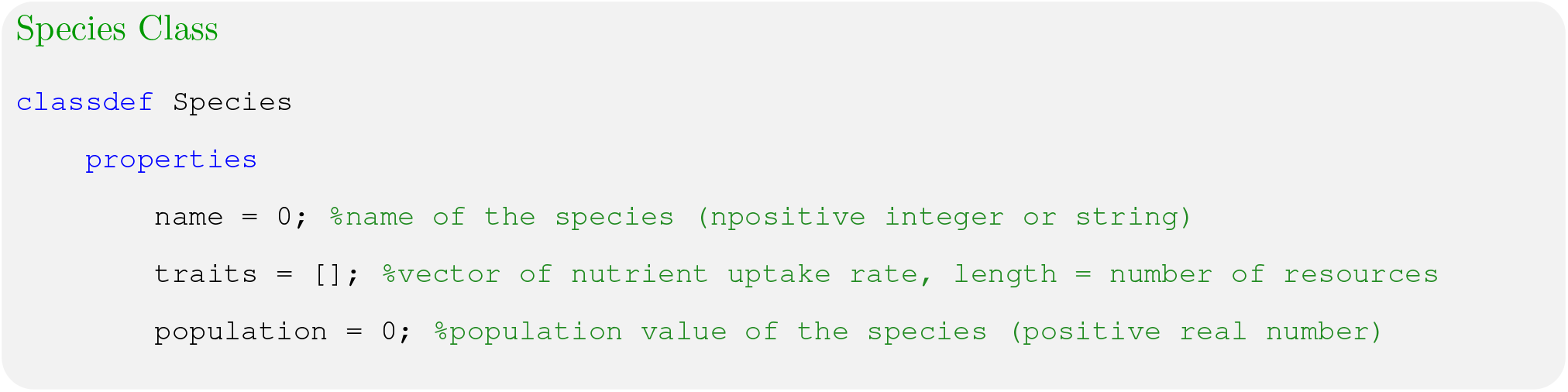

## name

Every species has a name, *σ* defined as a positive integer or as a string.

## traits

Every species is haracterised by certain traits, such as size, sedimentation rates, self-shading coefficients. With the resource-competition model described by dynamics.m only the metabolic strategies 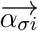 are considered. Additional traits an be incorporated in the traits array.

## population

population size, *n*_*σ*_ defined as a positive real number

## 7.2.3 Deme.m

A deme is a spatially isolated subcompartemnt of the ecosystem, where the the concentrations of resource are assumed to be homogeneous. A Deme is characterised by the following properties:

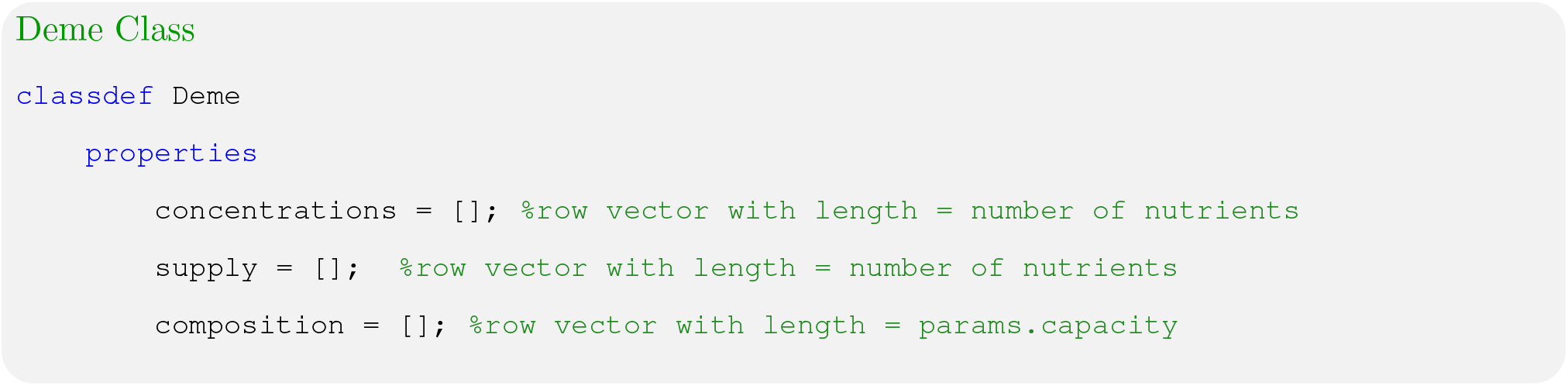

## nutrient concentrations

concentration of resources available 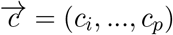

## nutrient supply

rates of nutrient supply, 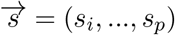

## species composition

list of Species objects, representing the composition of species, 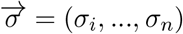 present in the given deme.

## 7.2.4 Plankton.m

The Plankton class defines the ecosystem as a matrix of demes.

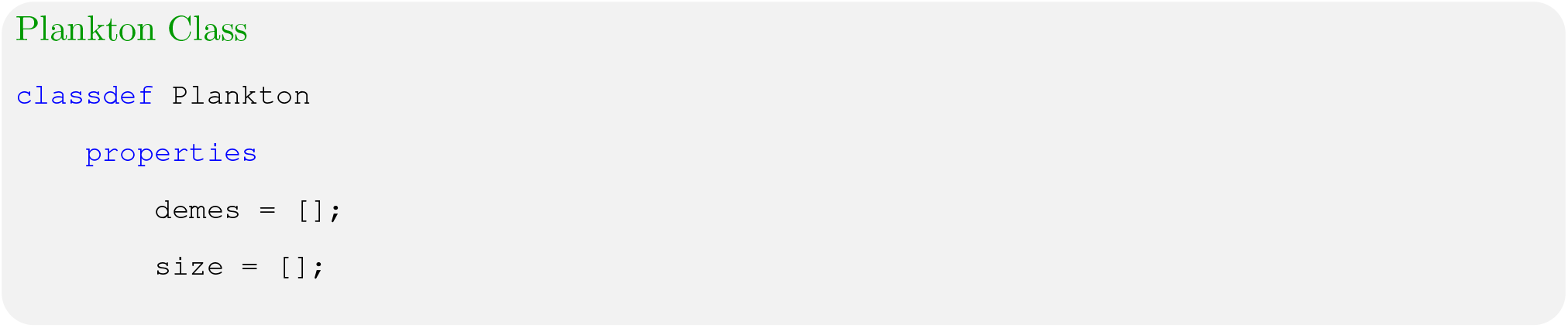

## demes

matrix of Demes objects

## size

dimensions of demes matrix

## 7.2.5 dynamics.m

The dynamics.m files defines the system of ordinary differential equations that solve the population and concentration dynamics in every deme. Every species is characterised by the coefficients of rate of utilisation for each nutrient *α*_*σ*_ = (*α*_*i*_, …*α*_*p*_). Biologically, *α*_*σi*_ is proportional to the number of enzyme molecules allocated by the organism to importing and processing *i* (Posfai et al. 2017). Enzymes for different nutrients may have different costs *w*_*i*_, but to account for “trade-offs,” Posfai et al. (2017) assumes that all organisms have the same fixed enzyme budget *E*

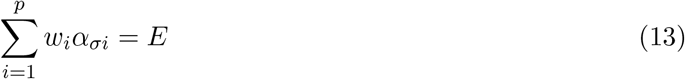

The nutrient uptake function, *r*_*i*_, is described by the monotone increasing Monod function, which saturates at high concentration of resource.

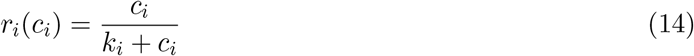

The growth-rate function of a cell *σ* is therefore given by the sum of the sum of the rates of the resource consumed (all nutrients is assumed to contribute to growth):

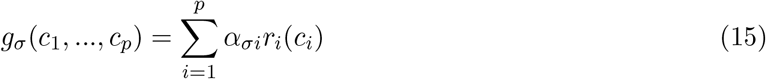

The final population dynamics for each species is given by a set of ODEs describing each species growth

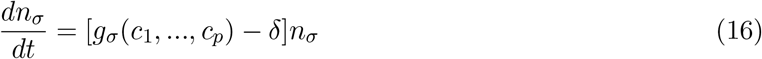

Each deme is assumed to be a well-mixed system such that the concentration of nutrients is homogeneous and is determined by the nutrient supply rates *s* = (*s*_1_, *…s_p_*), by the uptake of nutrients by organisms *r*_*i*_, and by a degradation or loss rate *μ*_*i*_. The kinetics of nutrient concentration *c*_*i*_ is equal to:

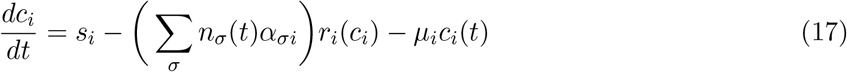

## 7.2.6 simulator.m

This is the main function of the plankton-simulator. It executes config.mat files generated by config.m and saves the outputs as config-data.mat The simulation algorithm is better explained with the wrokflow below:

1. Read config.mat files from “input” folder
2. Load parameters, initial species populations and resource concentrations
3. Seed every deme with a number of species specified by params.capacity
4. Assemble plankton according to dimensions set by params.size (
5. Check if the iteration if equal to the total simulation time
6. If not, the “simulate” function numerically solves the ODEs for species and resource dynamics using ode45s function (built in in MATLAB)
7. Execute “natural selection” function, which kills species with population size below value set by params.cutoff
8. Execute shuffle function, which draws a random species in every deme (weighted by population size) and appends a fraction (set by params.shuffle-amount) of its population into a random deme (inversely weighted by the Euclidean distance between the two demes)
9. Update each deme with population from invader species
10. Update total plankton simulation
11. If iteration = T, scripts stops and saves data as onfig-data.MAT files into “output” folder

**Figure 18:**
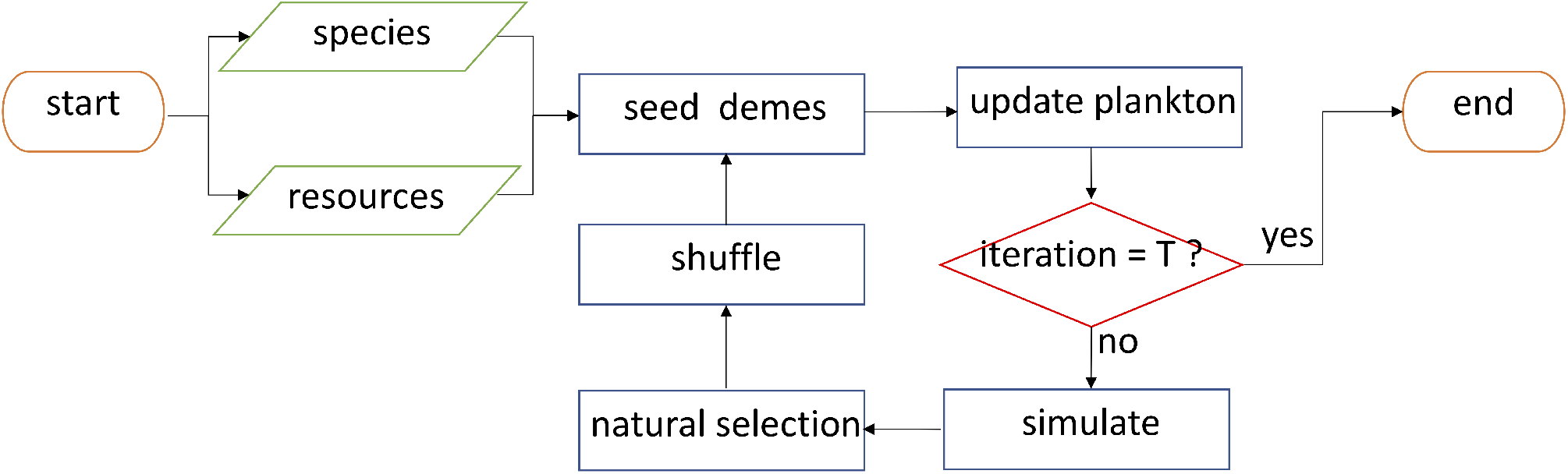
Stochastic Ecosystem Assembly - Algorithm

## 7.2.7 config.m

Scripts that allows user to specify parameters and simulation options and save them into “input” folder as config.mat files.

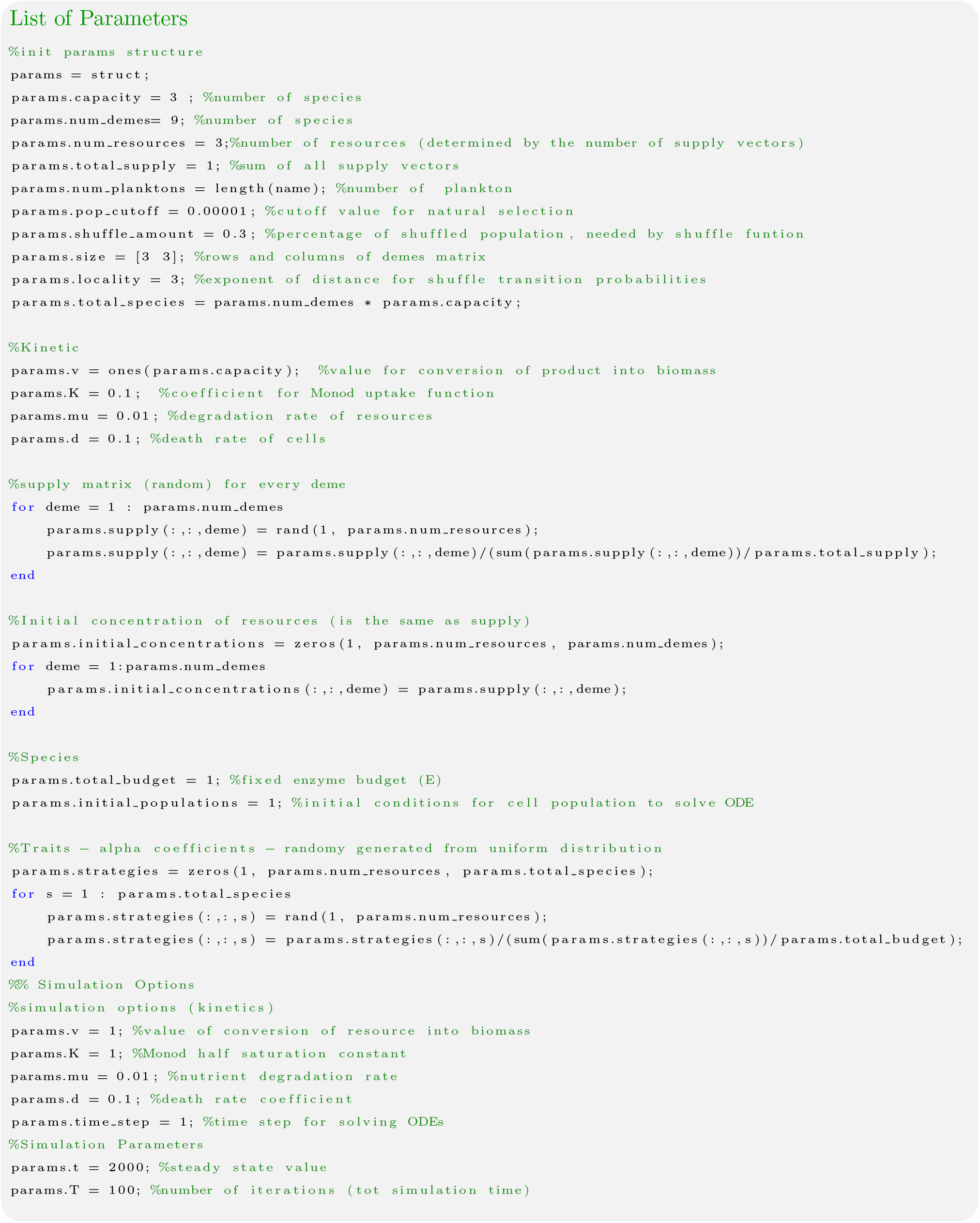

## 7.2.8 data reader.m

Script that analyses data and produces plot as shown in the main text.

## 7.3 Parameter Scanning

## 7.3.1 Steady State Time

## 7.3.2 Shuffle Locality

The shuffle probability is

**Figure 19:**
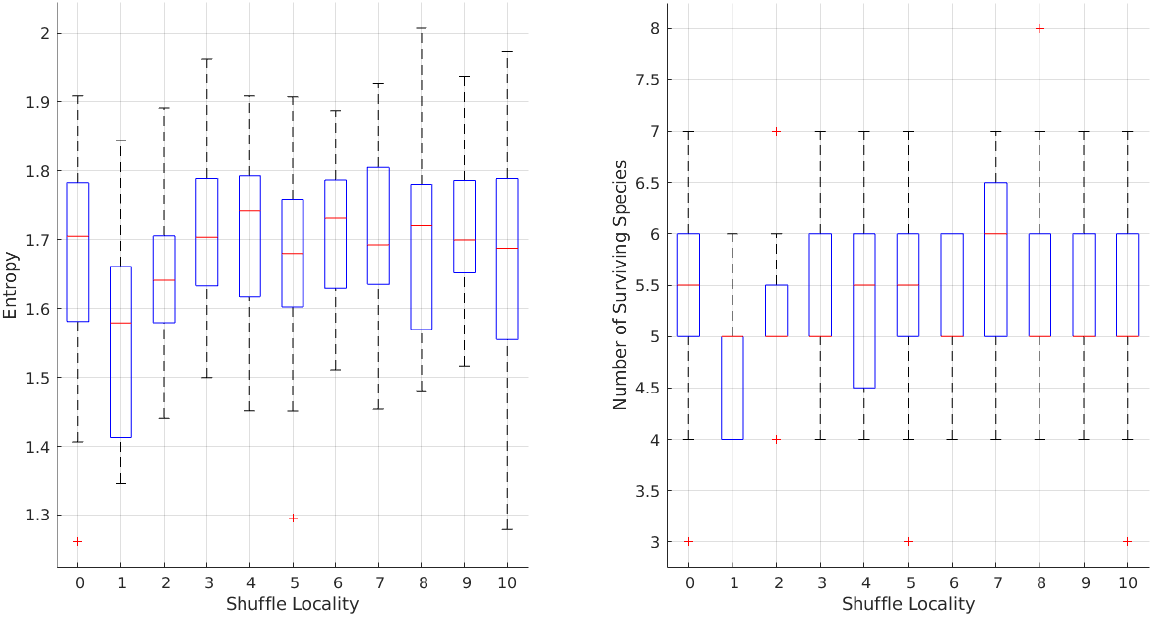
Entropy at different shuffle localities

**Figure 20:**
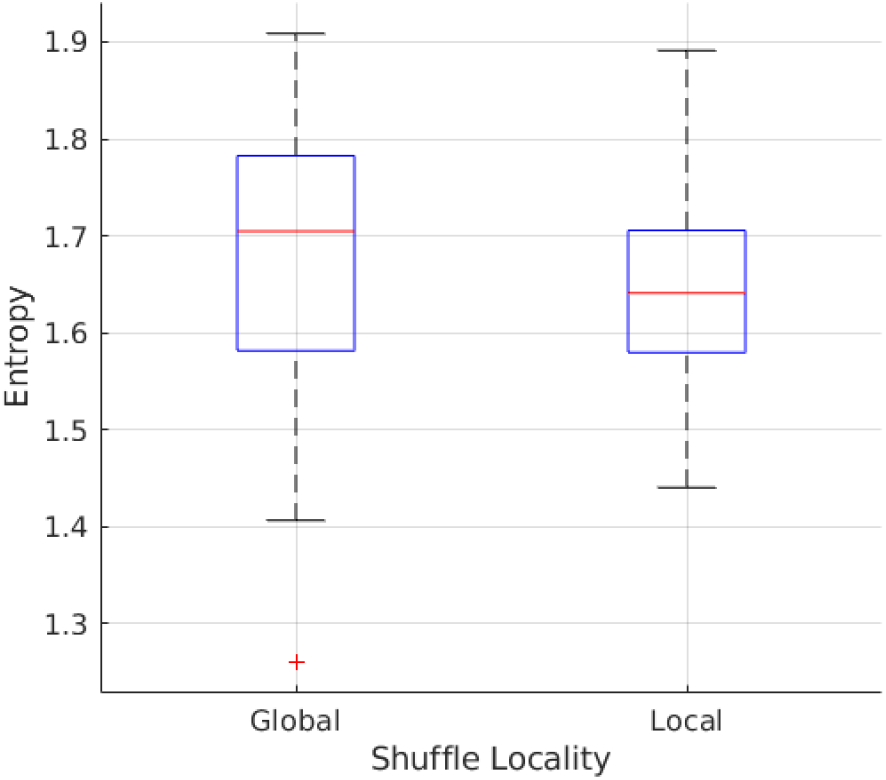
Entropy at different shuffle localities

**Figure 21:**
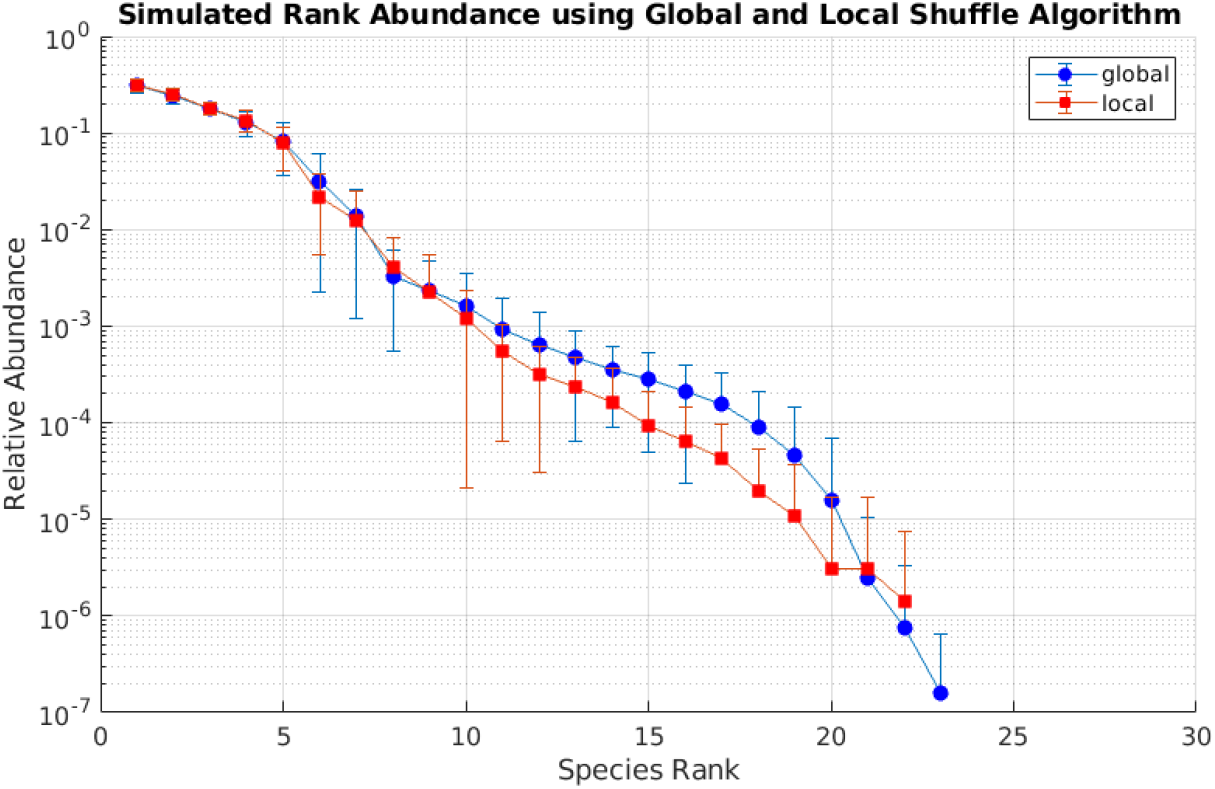
Rank Abundances Comparison, *l* = 0 and *l* = 2

## 7.3.3 Number of Species per Deme

To check if Shannon’s index is good biodiversity index a positive control simulation was run at different parameters of n: number of species per deme. Increasing the number of species seeded in each deme was expected to increase biodiversity index.

**Figure 22:**
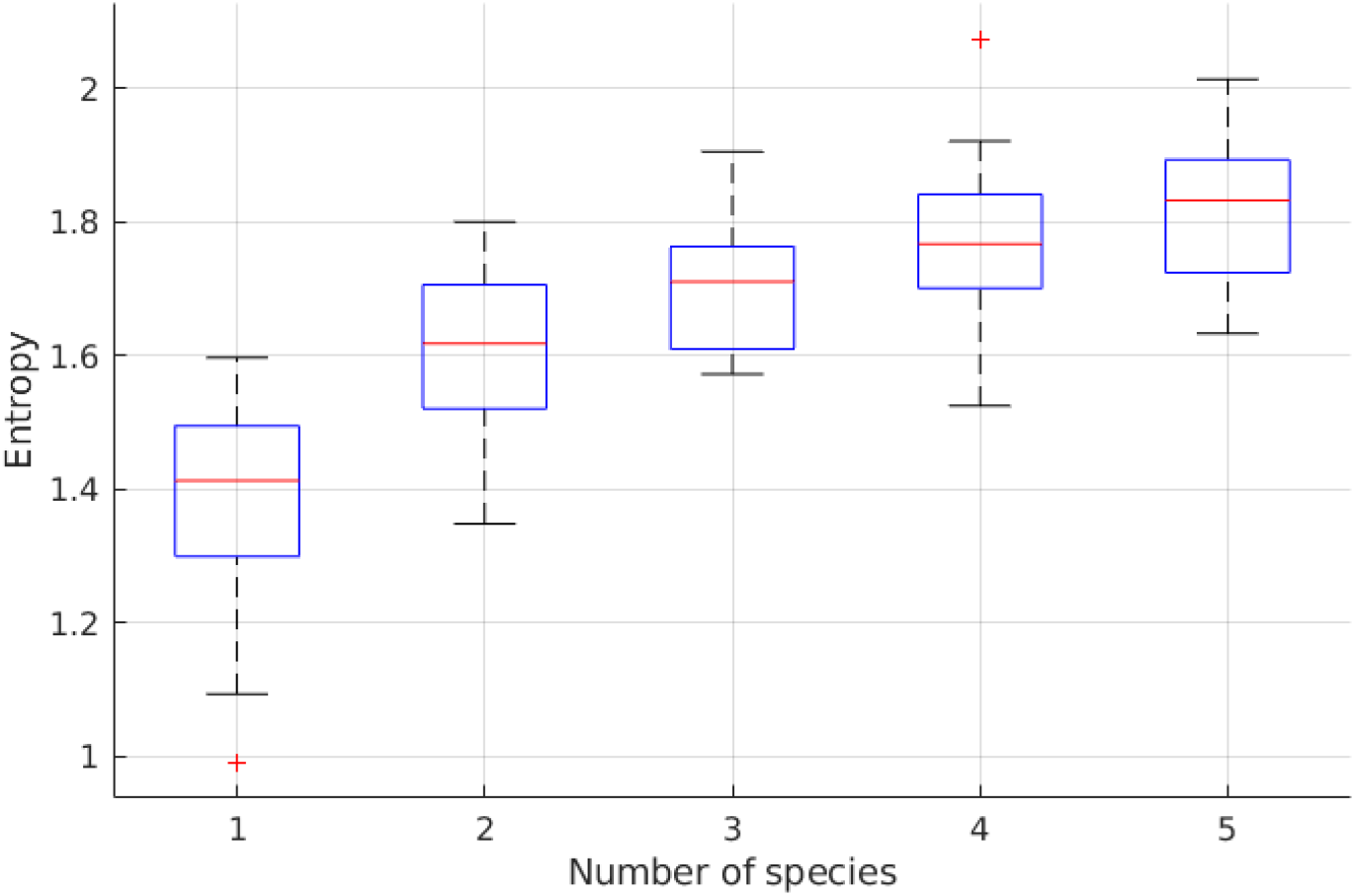
Ecosystem Shannon’s Entropy as a function of number of species per deme. Shown are boxplots from 20 observation per case, red line represents the mean values and the whiskers standard deviations.

Biodiversity index increases as the number of species per deme increase. It seems to saturate at high species per deme number. Probably because that is the maximum biodiversity for the given amount of nutrient.

## 7.4 Deviations in Fixed Enzyme Budget

**Figure 23:**
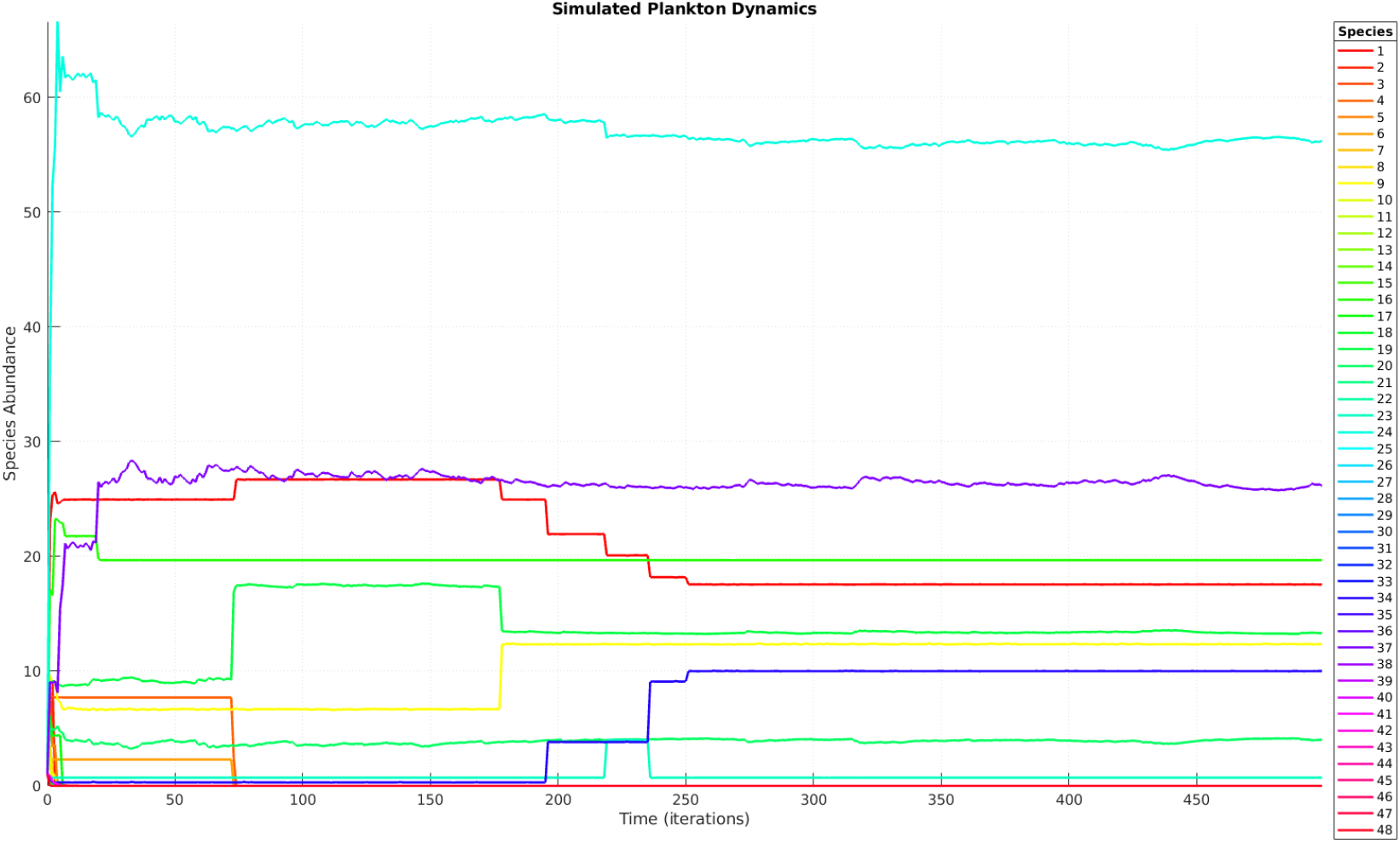
Metacommunity population dynamics with Σ = 0.05

**Figure 24:**
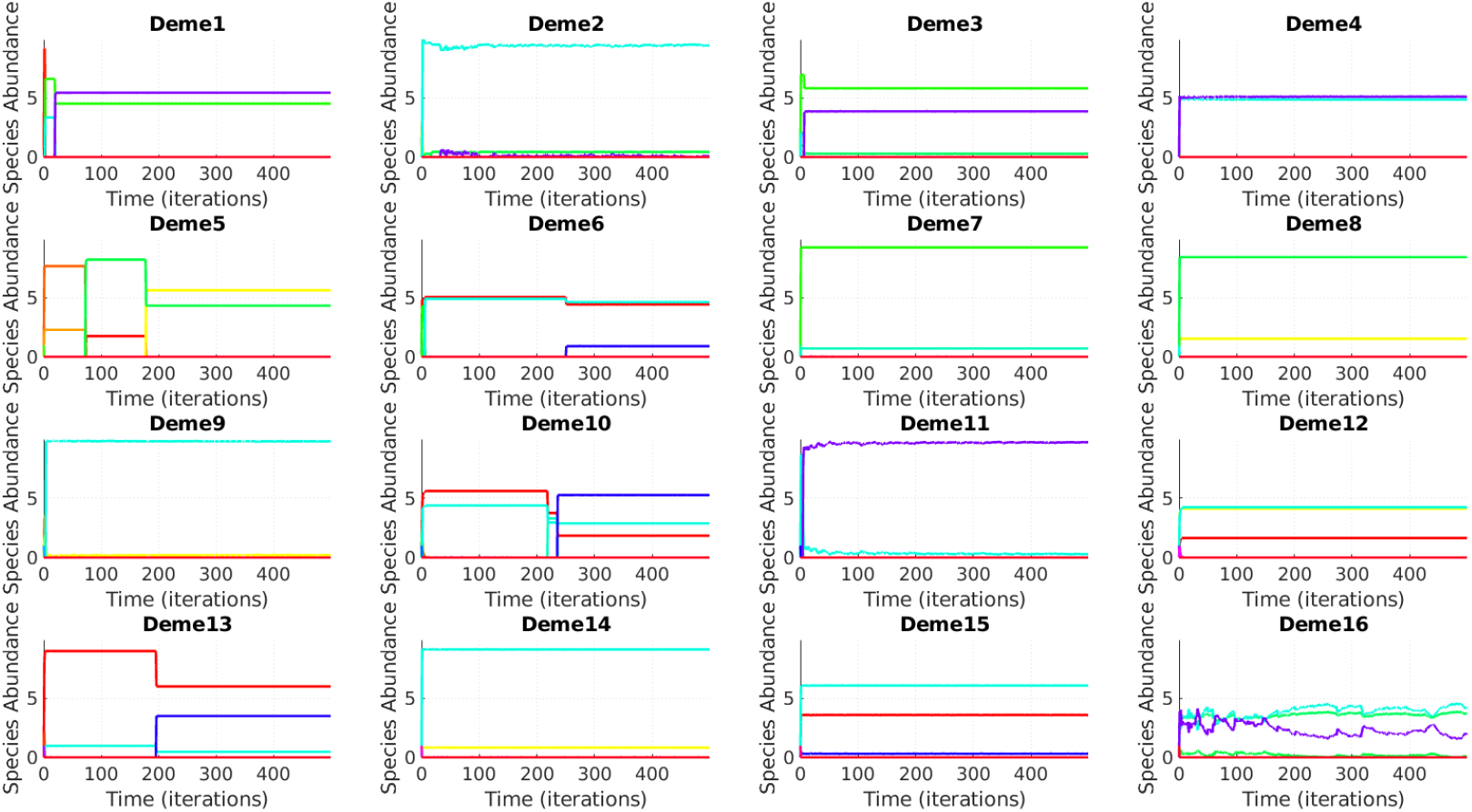
Community population dynamics with Σ = 0.05

**Figure 25:**
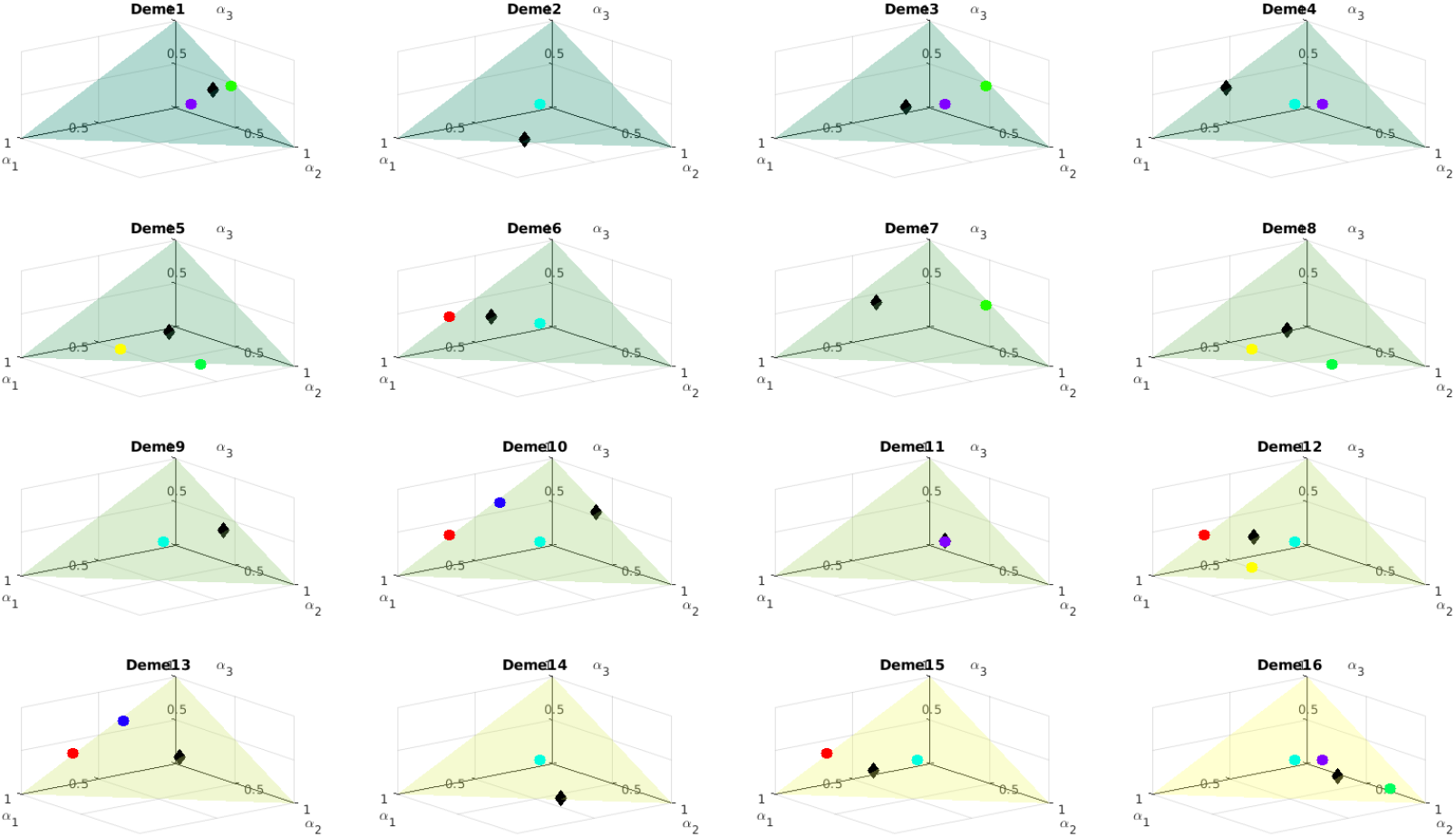
Space of resources utilisation at t=500 with Σ = 0.05

## Notes

https://github.com/scaralbi/plankton-simulator

### Competing Interest Statement

The authors have declared no competing interest.

